# Mutant p53 binds RNA to drive mitochondrial dysfunction

**DOI:** 10.1101/2025.11.12.688127

**Authors:** Wuyue Zhou, Alice Long, Cameron J. Douglas, Dalila Gallegos, James M. Burke, David W. C. MacMillan, Ezgi Hacisuleyman, Ciaran P. Seath

## Abstract

Tumor suppressor protein 53 (p53) is a transcription factor that is deregulated in 50% of cancers. Often termed the guardian of the genome, p53 is responsible for maintenance of genomic stability, cell cycle arrest, DNA repair, senescence, and apoptosis. In cancer cells, deregulation of p53 often occurs through mutations in the DNA binding domain which lead to a loss of the transcriptional activity. While 100’s of somatic mutations in the DNA binding domain are known, a small number of mutants are enriched in cancer, suggesting a gain-of-function role. Here we deploy an intein-based approach to localize µMap photoproximity labeling to p53 to define novel interactions contributing to the loss and gain of function roles of 5 separate hotspot mutants. These data revealed that G245S and R273H binds to RNA through its C-terminal domain. We show through CLIP experiments that mutant p53 has an RNA binding motif that conserved across mutants and is enriched in 3’UTRs, promoting ribosomal localization and labeling of proteins at the mitochondrial surface. We further demonstrate that the RNA binding ability of mutant p53 promotes altered miRNA processing and mitochondrial dysfunction providing mechanistic rationale for historically reported but poorly understood phenotypes.

## Introduction

The tumor suppressor protein p53 lies at the heart of cellular responses to stress. As a transcription factor, p53 maintains genomic integrity by activating a network of genes that coordinate DNA repair, cell cycle arrest, senescence, and apoptosis.^1,2^ This central role in protecting against oncogenic transformation has earned it the title “guardian of the genome.” p53 is one of the most frequently altered genes in human cancer, being mutated in almost 90% of all tumors.^1,3^ Somatic mutations that disrupt its DNA-binding domain (DBD) disable its transcriptional programs, abolishing tumor suppression and, in many cases, conferring new oncogenic properties (Fig. 1).^4^

**Fig. 1.**
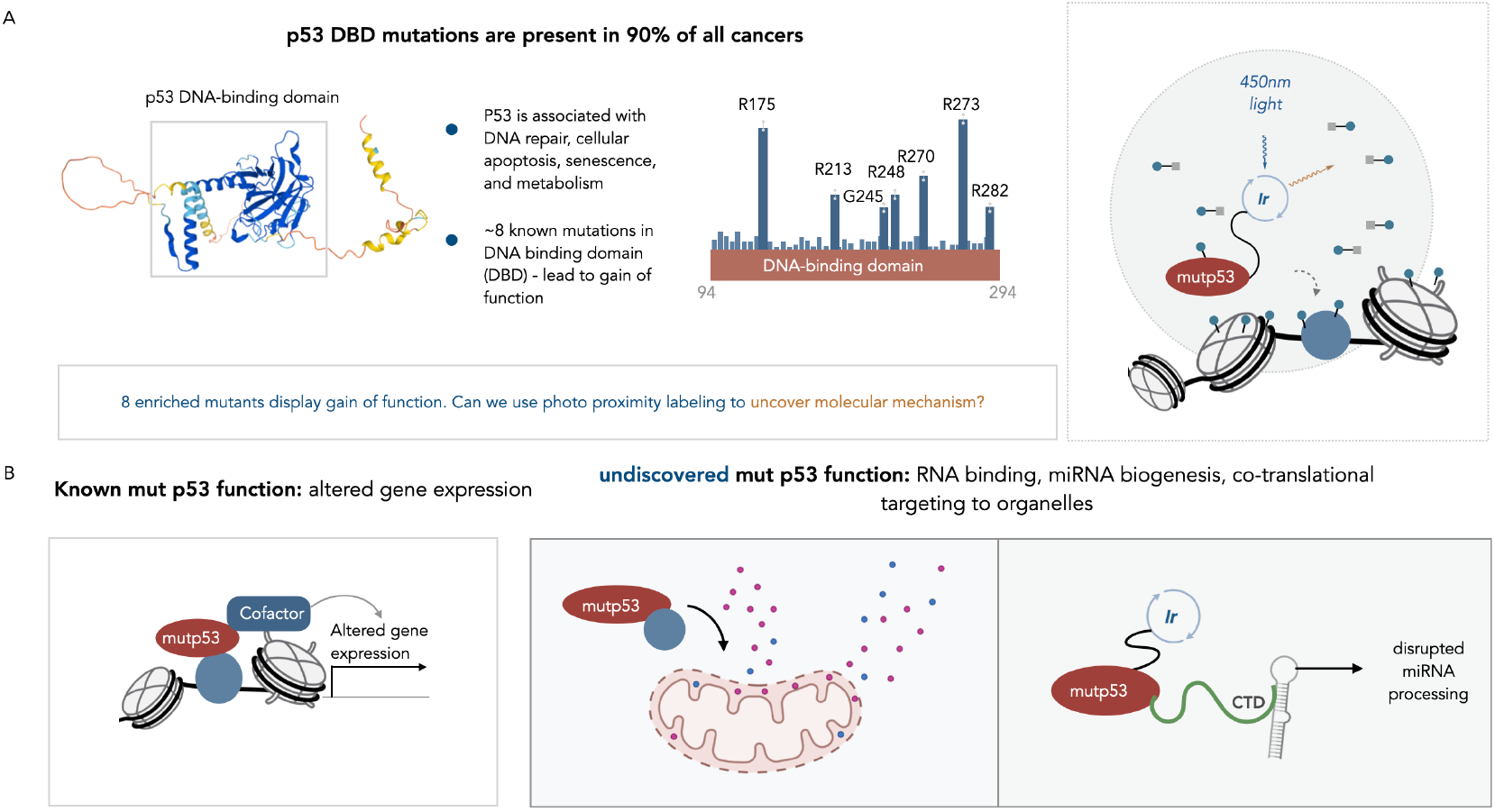
Unraveling mutant p53 function with µMap labeling. **(A)** somatic mutations in the DNA binding domain (DBD) in p53 are found in most cancers. A selection of these mutants is enriched in cancer and are termed “hotspot mutations”. In this study, we will use μMap proximity labeling to identify how these mutations change the protein-protein interactions around p53. **(B)** Hotspot mutants are known to lead to altered gene expression through mislocalization at genetic loci. μMap labeling reveals novel gain of function roles of mutant p53, including RNA binding, miRNA biogenesis, mitochondrial regulation, and co-translational targeting to organelles.

Among the hundreds of known p53 mutations, a small subset—often referred to as “hotspot” mutations—are recurrently enriched across tumor types and account for nearly three-quarters of all DBD variants.^3^ Their prevalence suggests that these mutants do more than simply lose transcriptional activity; rather, they may acquire distinct molecular functions that drive tumorigenesis.^3–8^ Uncovering the mechanisms behind these gain-of-function behaviors remains one of the central challenges in understanding mutant p53 biology.

Previous studies have used co-immunoprecipitation^7,9^ and chromatin immunoprecipitation^7^ to explore the interactions and genomic localization of mutant p53. These approaches have illuminated key binding partners and transcriptional programs but also have critical limitations. Co-IP methods capture stable protein–protein interactions but fail to preserve interactions mediated by nucleic acids or higher-order chromatin structures. Conversely, ChIP assays reveal DNA binding patterns but provide only indirect information about the dynamic and context-dependent complexes that define mutant p53 function.

To overcome these challenges, proximity labeling (PL) has emerged as a powerful approach to map molecular interactions in their native environments.^10–12^ PL methods can capture transient, weak, or nucleic acid–scaffolded interactions that are otherwise inaccessible to traditional biochemical techniques. Building on these advances, we sought to apply μMap^13^—a photocatalytic proximity labeling strategy developed for high-precision interactomics—to study how hotspot mutations reshape the molecular landscape surrounding p53. μMap operates through the short-lived generation of carbenes that label proximal biomolecules within nanometer-scale distances^14^, providing an exquisitely sensitive probe of local molecular architecture. This property makes μMap ideally suited to distinguishing subtle structural or conformational differences between wild-type and mutant proteins (Fig. 1, Fig. 2E).

**Fig. 2.**
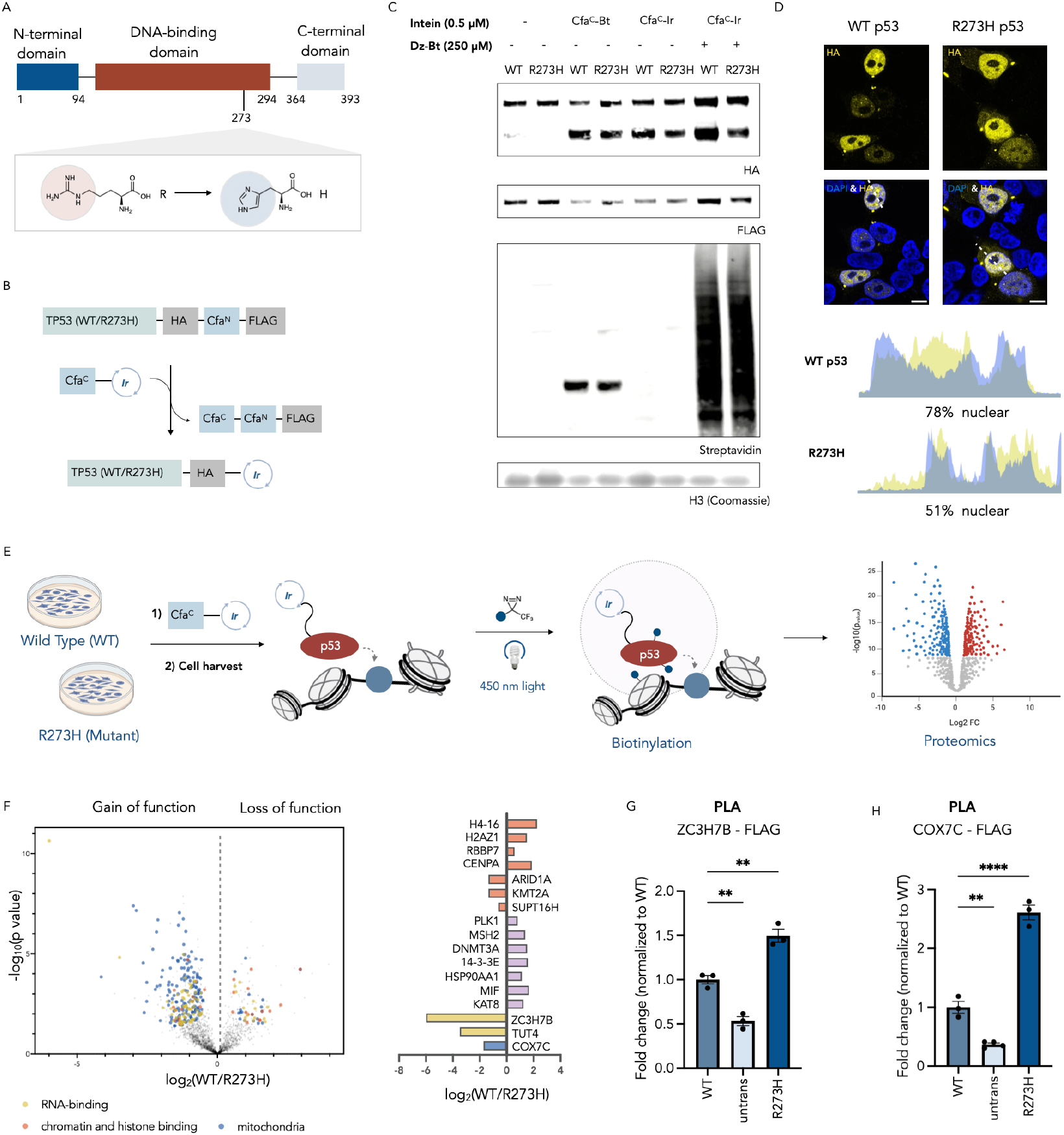
µMap proximity labeling uncovers the interactome of mutant p53. **(A)** Cartoon showing R273H mutation in the DBD of p53. **(B)** Scheme of intein splicing reaction. The N-terminal intein (Cfa^N^) is fused to p53, while the C-terminal intein (Cfa^C^) is conjugated to an Ir photocatalyst. Splicing leads to traceless conjugation of Ir onto p53. **(C)** Spliced product is only observed while treated with C-terminal intein (Cfa^C^). Streptavidin signal is only observed following treatment with Cfa^C^-Ir, Dz-Bt, and light. **(D)** Subcellular localization of wt p53 and R273H p53 in HEK293T cells. Wt p53 is exhibits a more concentrated nuclear localization (n≥5, two-sample t-test). Scale bar = 10 µm. **(E)** Scheme of µMap proximity labeling experiment. HEK293T cells transiently expressing mutant or wt p53 were harvested, treated with intein-conjugated Ir (Cfa^C^-Ir), and labeled with 450 nm light. Labeled proteins were enriched and subjected to proteomic analysis. **(F)** Interactome shift caused by p53 R273H mutation. Left panel: A volcano plot derived from a two-sided t-test comparing wt p53 interactors to R273H p53 interactors. FDR<0.05. RNA-binding, chromatin and histone binding, and mitochondrial proteins are annotated. Right panel: enrichment of selected proteins in the dataset. **(G)** Proximity ligation assay (PLA) confirming the interaction between p53 and ZC3H7B (n = 3, one-way ANOVA test followed by post hoc Dunnett’s multiple comparisons comparing to untransfected control. **P<0.01). PLA puncta per field of view were quantified for analysis. **(H)** Proximity ligation assay (PLA) confirming the interaction between p53 and COX7C (n = 3, one-way ANOVA test followed by post hoc Dunnett’s multiple comparisons comparing to untransfected control. **P<0.01, **** P<0.0001). PLA puncta per field of view were quantified for analysis.

To bring μMap labeling directly to p53 in its native nuclear context, we leveraged a split-intein delivery system recently developed in our laboratory^15,16^ (Fig. 2B). In this approach, p53 is expressed as a fusion with one half of a split intein, flanked by small epitope tags. Upon cell permeabilization, the complementary intein fragment bearing a pendant iridium photocatalyst is introduced. Rapid intein splicing seamlessly incorporates the photocatalyst onto p53, enabling precise labeling of its immediate molecular environment without disrupting native cellular architecture. This system allows μMap profiling in permeabilized cells with intact nuclei—conditions particularly suited to studying a transcription factor such as p53, whose function is deeply intertwined with chromatin structure and RNA interactions.

Together, this platform provides a powerful means to interrogate how specific p53 hotspot mutations alter molecular interactions. By combining proximity labeling with targeted intein-based delivery, we aim to define the mechanistic basis of mutant p53 gain and loss-of-function behavior and uncover new pathways that contribute to tumorigenesis.

### Method Development

We initiated our study by expressing plasmids containing either p53-HA-Cfa^N^-FLAG or the mutant R273H in HEK293T cells which are insensitive to p53 expression. We used the DNA binding domain mutation, R273H, as an exemplar due to the availability of prior gain of function studies and its abundance in a variety of tumor types (Fig. 2A). We validated intein-splicing of our constructs using Cfa^C^-Biotin and Cfa^C^-Ir (Fig. 2C).

Incubation with Cfa^C^-Biotin for 3 0 mins at 37 °C led to a shift in the construct mass as measured by western blot, in addition to a corresponding streptavidin signal at a mass indicative of full length p53, consistent with extein loss and incorporation of biotin at the C-terminus of p53. Cells treated with Cfa^C^-Ir were then irradiated at 450 nm for 60 seconds in the presence of diazirine-PEG3-Biotin (Dz-Bt) to induce protein labeling. Streptavidin western blot showed significant biotinylation only in the presence of the transfected construct, Dz-Bt, and light (Fig. 2C). We next demonstrated localization of wild type and mutant construct R273H via confocal microscopy, showing significant nuclear localization in both cases, with some non-nuclear signal observed in the mutant (Fig. 2D).

To ensure our construct can produce a high-quality interactome for p53, we performed proximity labeling experiment on wild-type p53 and an untransfected control. Each condition was performed with three biological replicates with two technical replicates per biological replicate to account for experimental variability. For hit-determination, we defined significance thresholds of log_2_FC>1 and detection in at least 4 positive replicates. Compartmental analysis of the identified proteins showed that 51% of all IDs were localized to the nucleus, 21% in the mitochondria, and 28% from other cellular compartments. Gratifyingly, we observed strong enrichment of p53 itself and several canonical binding proteins (*EP300*^17,18^, *KAT2A*^19^, *PML*^20,21^, *ATR*^22^) among our top hits, with enriched GO terms relevant to p53 function (Fig. S1A).

### p53 R273H blocks chromatin binding and associated interactions

Upon successful PL of p53, we proceeded to generate interactomics data for wild-type and R273H p53 via label free chemoproteomics to determine if we could delineate between interactomes of proteins that differ by only a single amino acid residue.

Analysis via volcano plot revealed 112 proteins significantly enriched in wild type samples (loss of function), and 337 were significantly enriched in mutant (gain of function) (Fig. 2F). GO analysis of the loss of function hits showed significant enrichment of chromosome organization, mitotic cell cycle, regulation of DNA and RNA metabolism, all key functions of wild-type p53^2,23–25^, consistent with a loss of DNA binding capacity and chromatin localization (Fig. S1B). This was supported by enrichment of 15 proteins related to histones, histone binding and chromatin organization (e.g., H4-16, H2AZ1, RBBP7, CENPA) in the wild-type interactome (Fig. 2F). We also observed a robust enrichment of proteins associated with DNA-related processes closely tied to p53 function including PLK1^26^, MSH2^27^, and DNMT3A^28^. In addition to chromatin localized proteins, several known proteins that regulate p53 function through direct interactions were enriched in the wild-type samples. For example, the 14-3-3ε, which interacts with the disordered C terminal domain of p53 and promotes tetramerization and transcriptional activity.^29^ It has been reported that HSP90AA1 binds to p53 and stabilizes the DNA-binding domain.^30^ Likewise, binding to MIF, which regulates p53 transcriptional activity in response to immune challenge^31^, is diminished in the R273H samples. In addition to proteins that directly regulate p53 stability and function, we identified KAT8, an acetyltransferase, which acetylates K120 of p53^32^ and mediates p53 based transcription independent apoptosis.

### p53 R273H displays gain of function interactions

Gene Ontology (GO) analysis of the gain of function interactions of p53 (R273H) revealed significant enrichment of terms related to RNA processing (e.g., ZC3H7B, TUT4), mitochondrial organization (COX7C), translation, and aerobic respiration, pointing to novel functions of the mutant p53 protein (Fig. S1B). Additionally, we observed a small number of gain-of-function interactions with chromatin bound proteins (ARID1A, KMT2A, SUPT16H), although these were significantly fewer than proteins associated with RNA or mitochondrial function, supporting a mechanism where R273H reduces affinity to chromatin and redirects p53 away from its canonical transcriptional interactome towards other functions. To ensure the validity of our novel hits, we performed proximity ligation assays^33^ to measure the abundance of two of our most enriched interactions (ZC3H7B, COX7C) in fixed cells. Using this technique, we measured a 1.5-fold increase in proximity between R273H and ZC3H7B and a 2.5-fold increase in proximity with COX7C when compared to wild type p53 (Fig. 2G, H, Fig. S1D).

Importantly, IP-MS of the same constructs was not instructive, with few differential interactions observed, suggesting that cellular localization, rather than new binding surfaces drive differential protein-protein interactions and providing further evidence that IP-MS protocols may be confounded by post-lysis changes to protein interactomes (Fig. S1C). Furthermore, comparison of enriched genes to the bulk proteome showed that enriched proteins were a result of proximity rather than changes in protein levels (Fig. S1C).

### p53 hotspot mutations display different localization and binding affinities

We next investigated a subset of other known hotspot mutations (R248Q, R282W, G245S, and R249S) to identify trends between the mutants and assess whether the gain and loss of function interactions are conserved (Fig. 1A). Prior to chemoproteomic characterization, we confirmed the efficiency of construct expression and splicing with Cfa^C^-biotin, demonstrating effective intein splicing across all constructs tested (Fig. S2A). Following this, we determined the cellular localization of each construct through cellular fractionation and confocal microscopy (Fig. 3A). These data clearly show significant differences between p53 localization dependent on the mutation under study, with all mutants exhibiting higher cytoplasmic localization (Fig. S2B).

**Fig. 3.**
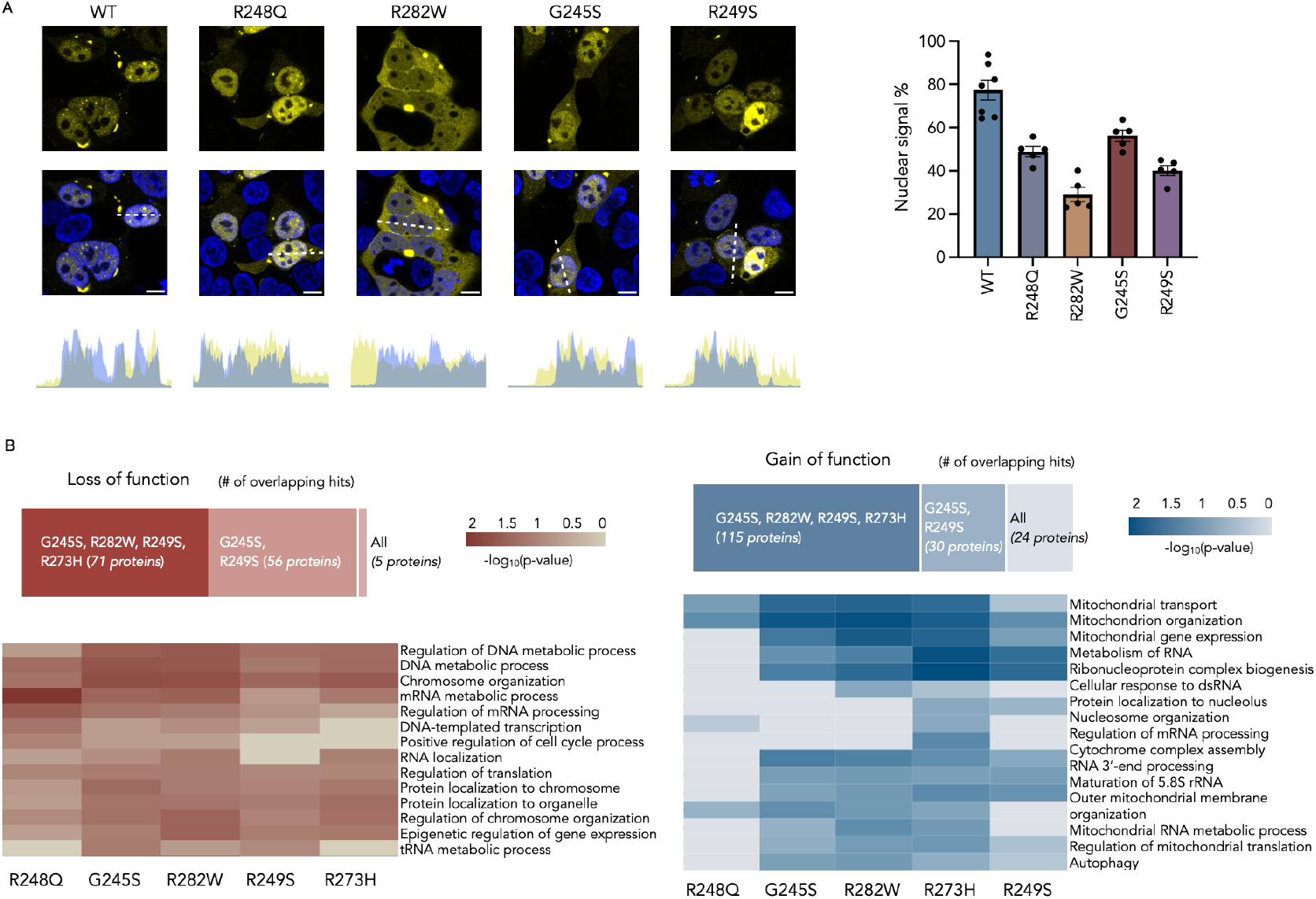
Unraveling functional changes in p53 with known hotspot mutations. **(A)** Subcellular localization of wt p53 and mutant p53 in HEK293T cells. Wt p53 is more highly localized in the nucleus than any mutant (n≥5, one-way ANOVA test followed by post hoc Dunnett’s multiple comparisons comparing to untransfected control. **P<0.01, **** P<0.0001). Scale bar = 10 µm. **(B)** Each mutant displays a unique interactome although some overlap is observed in loss of function, gain of function, and enriched gene ontologies. Hierarchical cluster analysis was performed based on the enrichment score of GO terms.

We next performed our proximity labeling experiment with these p53 hotspot mutants to determine shared gain-of-function roles between mutations. Their shared abundance across tumors of different lineage may suggest a conserved set of interactors that drive enrichment in cancer.^34^

Broadly we observe that the degree of interactome change is dependent of the identity of the mutant, and, consistent with their localization, that not all mutations bear identical interactomes. Hierarchical clustering of the data shows that the serine mutants G245S and R249S that alter p53 structure (*structural mutations*^35^) bear the most similarity and that R273H and R248Q, which block DNA binding directly (*contact mutations*^35^), also cluster together (Fig. 3B, Fig. S2D).^3^ Of the mutations tested, R248Q exhibited the weakest effect on the interactome, with 106 differentially enriched proteins, while R282W showed the strongest, with 1168 differentially enriched proteins (FDR=0.05) (Fig. S2C).

GO analysis of the gain-of-function terms across all five mutants showed consistent enrichment for mitochondrial localization (Fig. 3B). RNA processing was also significantly enriched in all mutants except R248Q. Together, these data suggest that somatic mutations in the p53 DNA binding domain promote mislocalization of the protein into the cytosol and mitochondria, enabling aberrant interactions with proteins involved in mitochondrial function and RNA processing. While each mutant displays unique features, several share significant overlap in their biochemical interactions (Fig. 3B). Based on these patterns, we selected R273H and G245S for further investigation, as they broadly represent the two major clusters and functional classes of hotspot mutations. Prior to follow up studies, we generated stable cell lines containing either wild-type p53, p53 R273H, or p53 G245S and measured the impact that the mutants had on the transcriptome through RNA-seq analysis. Pathway analysis showed that when compared to cells expressing wild type p53, both mutants effectively suppressed p53 based transcriptional activity, suggesting that they engage in dominant negative transcriptional behavior (Fig. S2E&F).

### Mut p53 binds RNA through the CTD and determines localization

Of the gain of function proteins associated with metabolism of RNA, we consistently enriched ZC3H7B, an RNA binding protein associated with miRNA biogenesis,^36,37^ in addition to several other RNA processing proteins (TUT4, TRA2B, SARNP, SRSF2) (Fig. 4A). As many of these proteins were enriched regardless of the identity of the mutant, we questioned whether in the absence of a high affinity DNA-binding interaction, mutant p53 exhibits altered localization and a preference for binding to RNA, and associated RNA binding proteins.

**Fig. 4.**
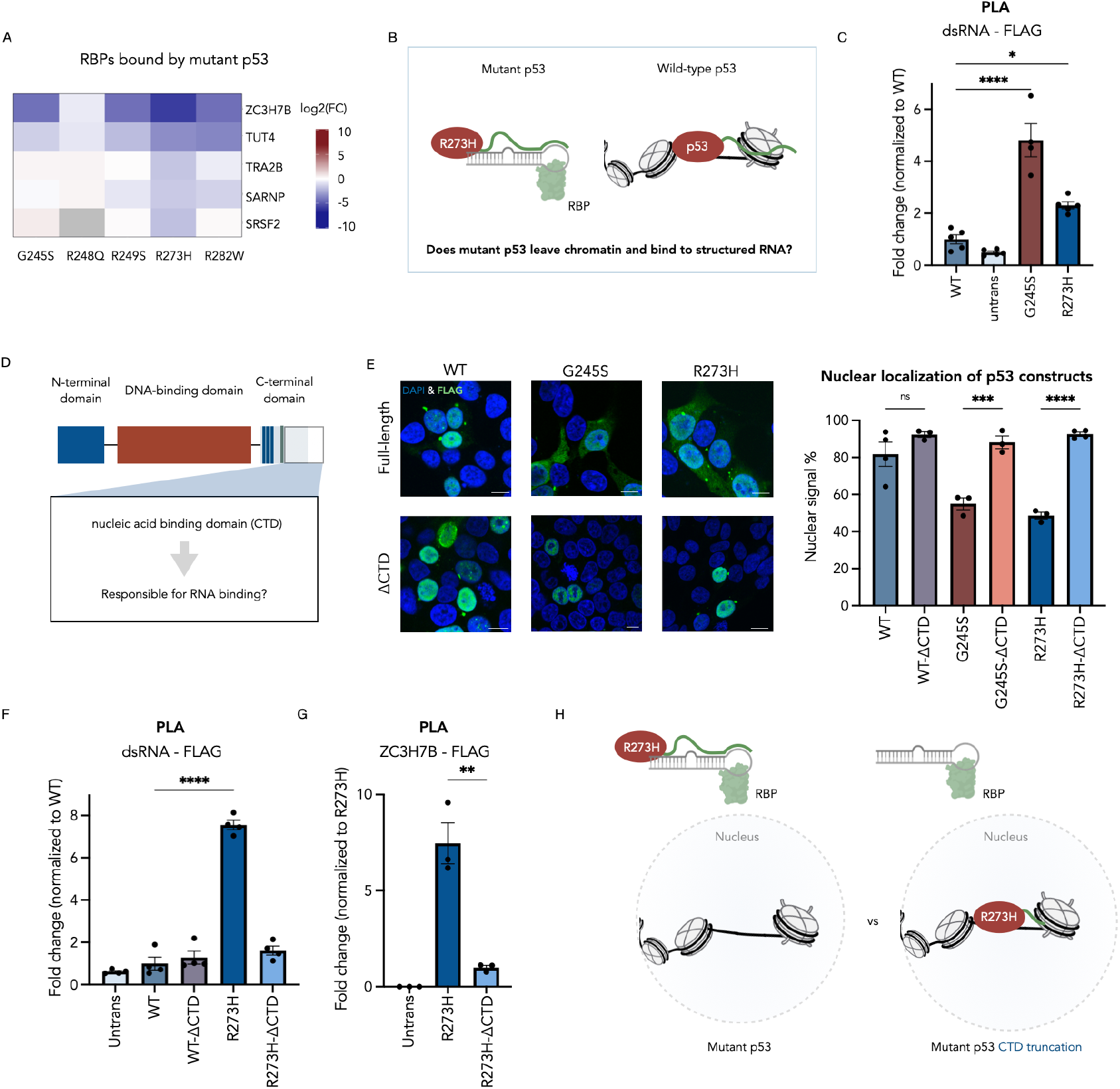
Mutant p53 binds RNA. **(A)** A Heatmap of RBP enrichment score across different mutations. **(B)** Cartoon illustrating how mutant p53 RNA binding might alter localization. **(C)** PLA confirmation of interaction between p53 and dsRNA (n = 3, one-way ANOVA test followed by post hoc Dunnett’s multiple comparisons comparing to untransfected control. *P<0.05, **** P<0.0001). **(D)** C-terminal domain has been previously reported as a non-specific nucleic acid binding domain. We question whether it is responsible for RNA binding in mutant p53. **(E)** Subcellular localization of full-length p53 and CTD-truncated p53 in HEK293T cells. Deletion of the CTD domain leads to nuclear localization in both mutant and wild type p53 (n≥3, one-way ANOVA test followed by post hoc Dunnett’s multiple comparisons comparing to corresponding full length control. ***P<0.001, **** P<0.0001). **(F)** PLA confirmation of the interaction between p53 and dsRNA after CTD truncation (n = 4, one-way ANOVA test followed by post hoc Dunnett’s multiple comparisons comparing to WT control. **** P<0.0001). **(G)** PLA confirmation of interaction between R273H p53 and ZC3H7B after CTD truncation (n = 3, two-sample t-test. ** P<0.01). **(H)** Cartoon showing R273H interaction with dsRNA and RBP. In full length R273H p53, CTD region binds to dsRNA and facilitates p53’s interaction with RBPs.

To test this, we performed PLA using antibodies against FLAG (WT, R273H, G245S) and an antibody that recognizes double-stranded and structured^38^ regions of RNA^39^ (*ds*-RNA K1) to quantify interactions between p53 and structured regions of RNA (Fig. 4B, C, Fig. S3A). We observed a 1.7-fold increase in interaction with *ds*-RNA for R273H and a 2.5-fold increase with G245S compared to wild-type p53, suggesting enhanced affinity of the mutant proteins for regions of double-stranded RNA *(>40 bp*). Broadly, RNA binding capacity by PLA correlated with RNA binding GO terms derived from proximity labeling (Fig. 3B). This result further supports the proposed model that mutant p53 may engage in non-canonical regulatory mechanisms involving RNA interactions rather than traditional DNA binding.

Given the complex structure of p53, we aimed to pinpoint the specific region of p53 that mediates its binding to RNA. We analyzed the RNA-binding capacity of mutant p53 (R273H) through domain-specific truncation, specifically targeting the C-terminal domain (CTD) (Fig. 4D). The CTD has been previously described as a non-specific nucleic acid binding domain^40^, suggesting it may be responsible for RNA localization, although it has been previously proposed to direct p53 to specific regions of chromatin^40,41^. To assess the impact of this truncation, we performed immunofluorescence staining to evaluate the subcellular localization of R273H, G245S and CTD-truncated variants. In cells expressing R273H, we observed that approximately 50% of mutant p53 localized to the nucleus, with a notable portion remaining in the cytoplasm—consistent with previous findings^42,43^ (Fig. 4E). Strikingly, upon truncation of the CTD, the nuclear localization of mutant p53 increased significantly, with approximately 90% of the protein now residing in the nucleus. Similar trend was observed for mutant G245S, of which nuclear localization increased from 55% to 88% upon CTD truncation (Fig. 4E). This shift in localization suggests that the CTD plays a crucial role in mediating the cytoplasmic retention of mutant p53^44–46^. We validated the role of the C-terminal domain (CTD) in RNA binding using PLA, observing a five-fold increase in RNA binding signal for full-length R273H compared to wild-type p53, wild-type ΔCTD, and R273H ΔCTD constructs (Fig. 4F, Fig. S3B). This significant difference strongly suggests that the CTD of mutant p53 is essential for its interaction with *ds*-RNA. To further support this observation, we performed PLA using ZC3H7B and FLAG-tagged R273H with and without a CTD deletion (Fig. 4G, Fig. S3C). We observed a notable increase in interaction between ZC3H7B and the full-length R273H mutant compared to both the CTD-truncated mutant and untransfected cells. These findings reinforce the notion that the CTD of mutant p53 facilitates its interaction with RNA-associated proteins, such as ZC3H7B, in addition to dsRNA (Fig. 4H).

Our above data indicate that mutant p53 binds structured RNAs (dsRNA regions). Notably, the biogenesis of miRNAs requires processing of longer pri-miRNA and pre-miRNAs, which are ~33-bp and ~22-bp in length, respectively. Thus, mutant p53 could potentially interact with these structures and alter miRNA biogenesis. To assess the functional impact of mutp53 binding to these miRNA precursor structures, as well as the proteins known to be involved in miRNA processing (ZC3H7B^37^, TUT4^47^), we performed miRNA-seq on cells stably expressing WT-p53, R273H-p53, and G245S-p53 (Fig. S4A, B). To differentiate between miRNAs that are regulated by the transcriptional or RNA-binding functions of p53, we filtered the data to identify RNAs that are selectively regulated at the mature transcript but remain unchanged at the precursor level (Fig. 5A). For G245S, we observed 42 differentially regulated miRNAs, of which 30 were only regulated at the mature miRNA level (P_adj_<0.1), while for R273H 26 out of 32 transcripts were regulated at the mature miRNA level, consistent with mutant p53-driven disruption of miRNA processing^25,48,49^ (Fig. 5B, C) Of specific interest were a set of tumor-suppressing miRNAs (miR-28^50^, miR-139^51^, miR-181^52,53^, miR-204^54^, miR-671^55^, and miR-885^56,57^) that were significantly regulated only at the mature form, suggesting that mutant p53 obstructs processing of their precursor hairpins (Fig. 5C, D).

**Fig. 5.**
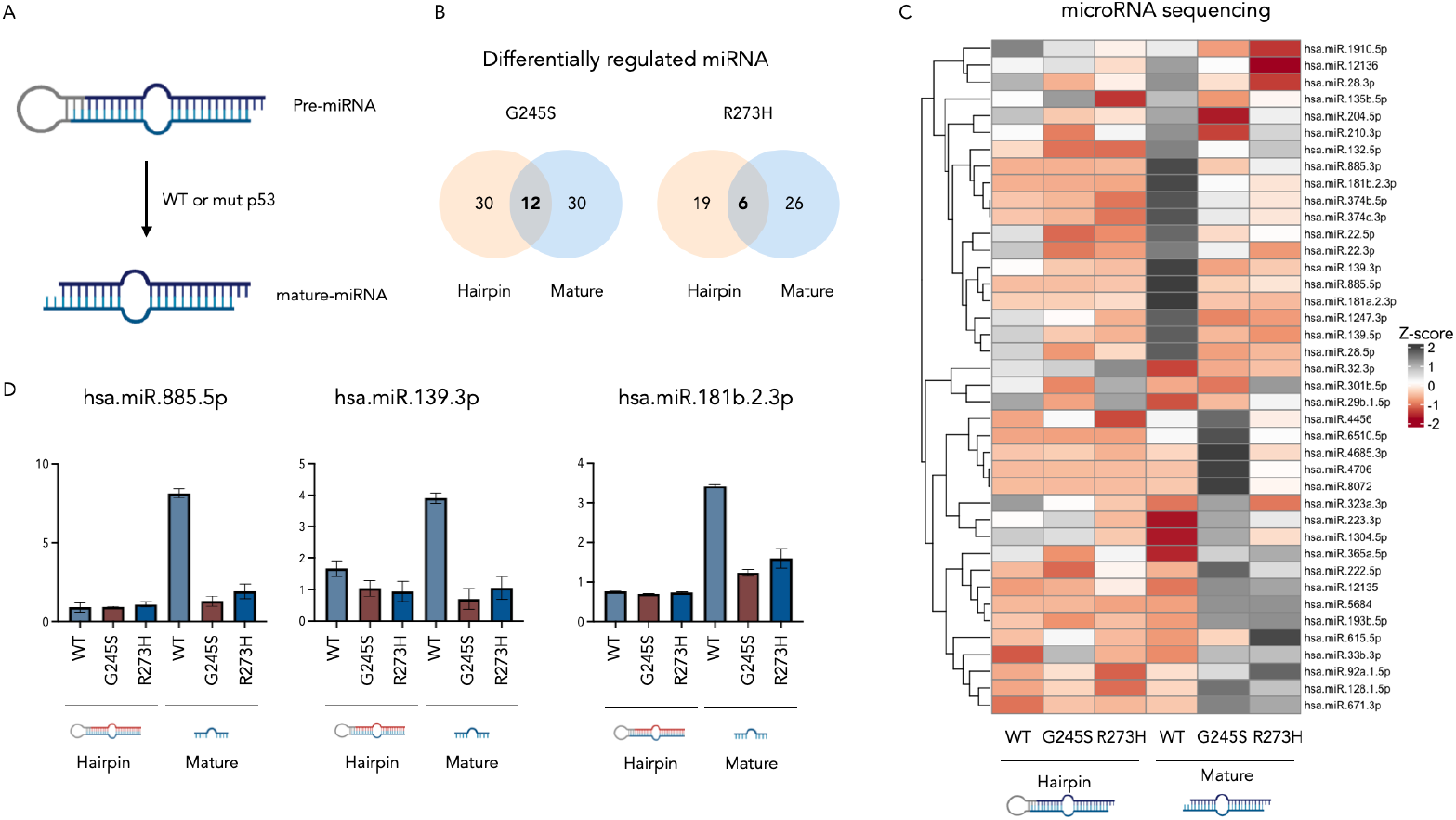
Mutant p53 regulates miRNA biogenesis. **(A)** A diagram showing p53 participation in miRNA processing. **(B)** Venn diagram showing differentially regulated hairpin and mature miRNA in mutant p53 (normalized to WT p53). **(C)** Heatmap showing z-score of miRNAs that are differentially regulated on the mature RNA level but not regulated on the hairpin level in both mutants. Z-score is calculated within each miRNA in both hairpin and mature RNA. **(D)** Individual bar plots showing selected miRNAs relevant to tumor suppression that are regulated at the hairpin level.

The selective regulation of mature, but not precursor, miRNAs suggests that mutant p53 disrupts RNA processing rather than transcriptional regulation. Together with our previous observations of increased dsRNA interactions and enhanced cytosolic localization, these findings point to a broader alteration in RNA-binding behavior. To directly examine this, we performed CLIP-seq on wild-type p53, R273H, and G245S (Fig. 6A) to define their RNA-binding profiles genome-wide. We observed significantly increased RNA-binding in both mutants, in line with our PL and PLA datasets (Fig. S5A). Each mutant bore a distinct binding profile to wild type as visualized by PCA, with good clustering among replicates (Fig. S5B). DESeq analysis showed a dramatic difference in transcript identity, with WT-p53 binding to intronic RNA and promoter regions, consistent with its nuclear localization. Both mutants demonstrated increased binding to 3’UTRs, with the most dramatic effect being observed in G245S (Fig. 6B), consistent with GO terms (RNA 3’-end processing) observed in our proximity proteomics (Fig. 3B). Furthermore, motif analysis showed that wild type and mutants shared enrichment for a common consensus sequence centered on GGCUGG, suggesting that p53 has an intrinsic RNA-binding preference (Fig. 6C). This motif was further enriched in the mutants, indicating that disruption of the DNA-binding domain enhances, rather than redirects, this RNA interaction, independent of the specific somatic mutation and consistent with our CTD-deletion findings (Fig. 6C, see Fig. S5C for additional enriched motifs). GSEA of bound RNAs across both mutations showed binding to ribosomal transcripts and those associated with co-translational targeting to organelle membranes, implying that mutant p53 binds to 3’UTRs of RNAs translated at ER^58^ and mitochondria^59^ (Fig. 6D). Re-analysis of our proximity labeling data supported this with enrichment of mitochondrial outer membrane proteins TOMM7, TOMM22, TOMM20 and TOMM40, in addition to co-translational chaperones AKAP1 and AKAP6 (Fig. 6E) that are previously found to be localized to the mitochondrial surface^60^.

**Fig. 6.**
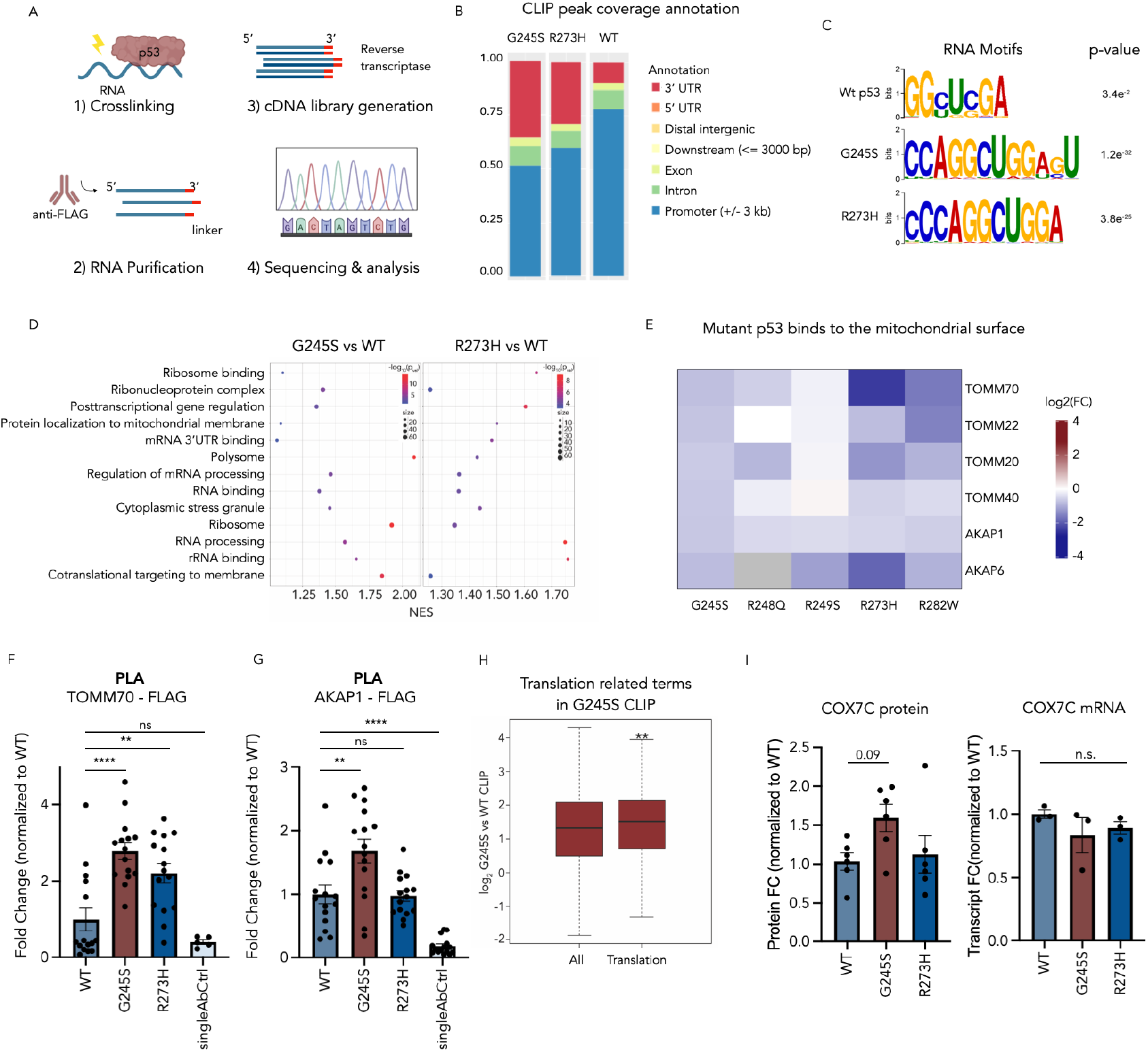
Mutant p53 binds RNA at mitochondrial surface. **(A)** Schematic of CLIP-seq workflow. **(B)** CLIP-seq peak coverage in genomic regions. **(C)** A consensus RNA-binding motif containing a common GGCUUG sequence is shared between wild-type and mutant p53. This motif represents the most enriched sequence in RNAs bound by wt and G245S p53 and the second most enriched motif in R273H p53. Motifs were identified using STREME analysis (MEME Suite) from peaks present in both biological replicates. (D) Normalized Enrichment Score (NES) from Gene Set Enrichment Analysis (GSEA) of CLIP-seq peaks. Gene sets commonly enriched in both mutant versus wild-type conditions are shown. **(E)** A heatmap showing enriched proteins that localize to the mitochondrial outer membrane identified through µMap PL across different mutations. **(F)** PLA confirmation of interaction between p53 and TOMM70 (n=15, one-way ANOVA test followed by post hoc Dunnett’s multiple comparisons comparing to WT control. **** P<0.0001). **(G)** PLA confirmation of interaction between p53 and AKAP1 (n=15, one-way ANOVA test followed by post hoc Dunnett’s multiple comparisons comparing to WT control. **** P<0.0001). **(H)** Among G245S-bound transcripts, RNAs associated with translation highly enriched. Translation-related proteins were retrieved from MSigDB (GOBP: Translation; GO:0006412). Statistical significance was calculated using the Wilcoxon signed-rank test. **(I)** Mutant p53-bound COX7C has increased expression but no upregulation at the transcript level. Left: COX7C protein expression level in global proteomics. Right: *COX7C* transcript level assessed by RT-qPCR (n=3, one-way ANOVA test followed by post hoc Dunnett’s multiple comparisons comparing to WT control.).

Consistently, PLA assays confirmed mutant-specific interactions between p53 and mitochondrial components (TOMM70, Fig 6F) as well as AKAP1 (Fig. 6G), supporting a model in which mutant p53 engages RNA and protein partners at the mitochondrial surface (Fig. S5D).

We next cross analyzed our CLIP and PL datasets to identify mitochondrial proteins whose transcripts are bound by mutant p53 and whose encoded proteins co-localize with mutant p53, potentially indicating a role for mutant p53 in co-translational targeting to the mitochondria. Among these, we identified that COX7C, an essential component of the electron transport chain^61^ was bound more by mutant p53 at both the RNA and protein levels (Fig. S5F). RT-qPCR analysis showed that the *Cox7c* transcript remained unchanged in all conditions, while the presence of mut-p53 increased its protein levels (as shown by global proteomics analysis) (Fig. 6I). We also noted several global trends that were supportive of our hypothesis. Correlation of the global proteomic and CLIP datasets revealed that translation-associated proteins not only showed increased abundance in the mutant proteome but also had their mRNAs bound by mutant p53 (G245S) (Fig. 6H). Furthermore, mitochondrial proteins, in addition to being enriched in the proximity labeling dataset of mutant p53, are also enriched in the global proteomes for both mutants studied (Fig. S5E).

Taken together, these data point to a coordinated shift in RNA and protein interactions that positions mutant p53 at the interface of mitochondrial and translation regulation, hinting at broader consequences for cellular metabolism.

### Mut-p53 leads to changes in mitochondrial function

A known function of wild type p53 is to translocate to the mitochondrial membrane leading to increased membrane permeabilization and cytochrome release.^59,62^ Our combined ‘omics suggests that this function exhibited by WT p53 may be enhanced upon loss of DNA binding capacity in the mutant protein.

To quantify the impact of mut p53 on mitochondrial function we measured mitochondrial mass, membrane potential, and ROS via MitoTracker microscopy assays. We found that mitochondrial mass almost doubled (MitoGreen) and mitochondrial potential (MitoRed) increased significantly in cells stably expressing R273H p53 and G245S p53 when compared to wild type, suggesting that mitochondrially localized p53 has a negative functional impact of mitochondrial health (Fig. 7A, Fig. S6A). We also investigated cellular reactive oxygen species and respiration. MitoTracker orange, which requires ROS to activate it prior to mitochondrial accumulation, showed a 1.3-fold increase in mut p53 compared to wild type (Fig. 7A, Fig. S6A). This data was confirmed by SeaHorse analysis which showed two-fold increase in basal and max respiration and ATP production in cells stably expressing mutant p53 when compared to wild type (Fig. 7B,C). Thus, mutant p53’s mislocalization and RNA-binding properties extend beyond molecular interactions to directly alter mitochondrial physiology (Fig. 7D).

**Fig. 7.**
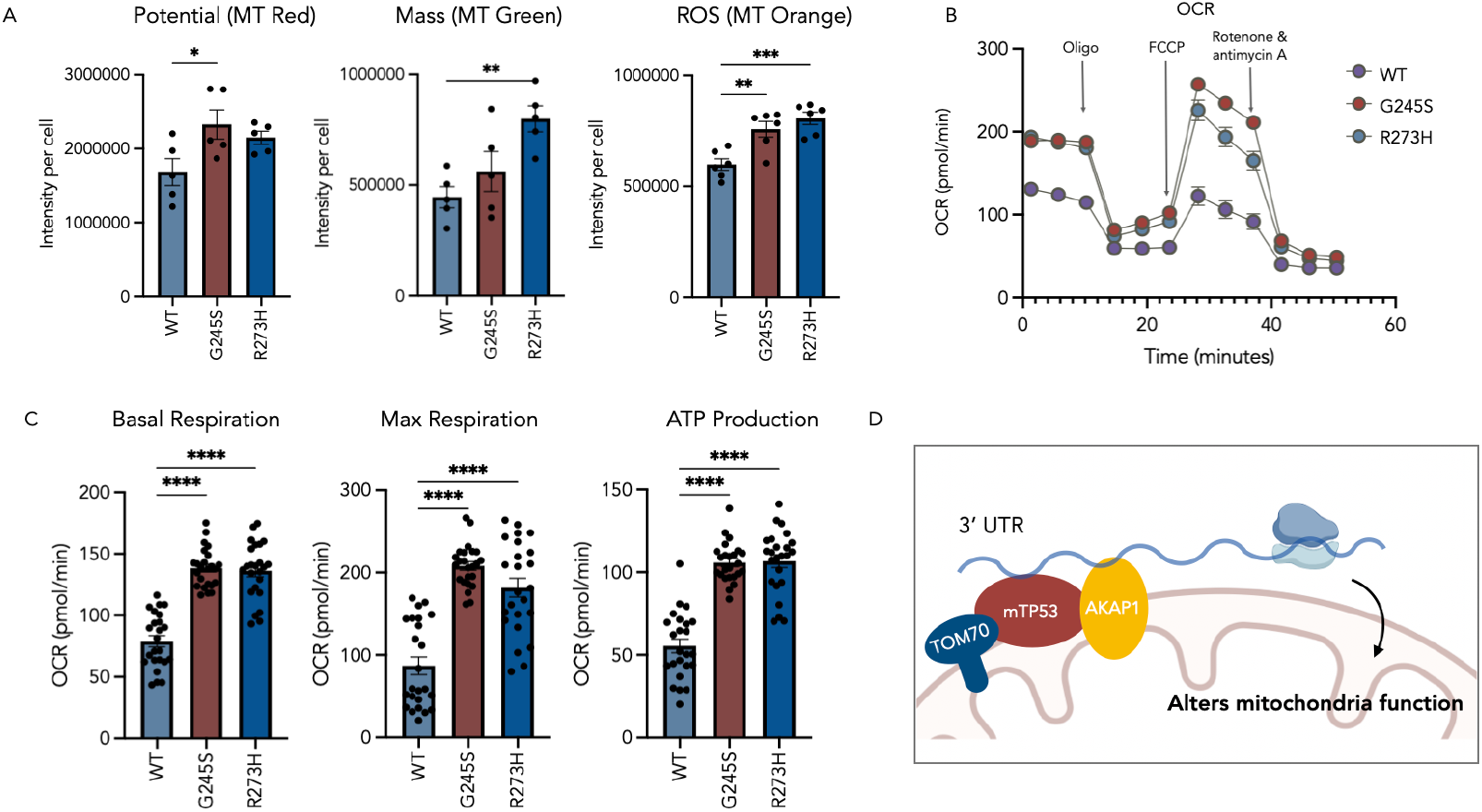
Mutant p53 alters mitochondrial function. **(A)** Expression of mutant p53 leads to changes in mitochondrial mass, membrane potential, and ROS level. (n≥5, one-way ANOVA test followed by post hoc Dunnett’s multiple comparisons comparing to WT control. **** P<0.0001). **(B)** Mitochondrial respiration was measured by Seahorse assay. **(C)** Basal respiration, max respiration, and ATP production was altered due to mutant p53 expression in cells. **(D)** A cartoon showing model of mutant p53 localizes on mitochondrial outer membrane and binds to RNA and AKAP1.

## Discussion

Mutant p53 is common feature across cancers with a strong genetic selection for a small number of somatic mutants in the DNA binding domain. The role of these mutations is a topic of much debate in the literature with strong recent evidence being presented that in mature cancer cell lines containing only the mutant (loss of heterozygosity), no phenotypic difference was observed between when the mutant protein was knocked out using CRISPR-Cas9. Despite this, the strong selective pressure for a small subset of mutants suggests they likely play a role in another aspect of carcinogenesis, such as oncogenic transformation or loss of heterozygosity.

In this study, we describe a biochemical analysis of the varied interactions exhibited by both wild-type and mutant p53, and how hotspot mutations redirect p53 localization and function. Our data clearly shows that these mutations promote certain situational functions of p53, such as modulating mitochondrial stress and RNA processing. While these functions are part of the complex repertoire of wild type p53 phenotypes, they appear to be overrepresented when the protein can no longer bind to DNA. We posit that once the strong DNA binding affinity of wild type p53 is lost or diminished, the non-specific nucleic acid binding domain in the CTD begins to dominate function, promoting protein translocation and RNA binding. While it remains unclear how these particular interactions could promote oncogenic transformation at the early stages of cancer, our datasets provide a biochemical basis for such investigations and provide commonalities between different hotspot mutations.

Despite these novel insights, our study contains several limitations. First, the intein based method we employ in this study requires the use of permeabilized cells, which will disrupt the cellular environment and will struggle to capture cytosolic interactions, leaving a gap in our knowledge. Second, we perform these experiments in a non-cancerous model cell line, HEK293T, which suppresses its own copy of p53. While this allows for the expression of all variants of p53 in a single controlled system, it does not capture the unique environment that each mutation is typically found in. Future work will focus on exploring mut p53 as an RBP in representative cancer models.

## Supplementary Information

Supplementary information is linked to the online version of the paper.

## Data Availability

All relevant data are included in the manuscript and supplementary information. Mass spectrometry data files have been uploaded to the MassIVE proteomics database (MSV000099500).

## Acknowledgements

Research reported in this publication was supported by the Office of The Director, of the National Institutes of Health under Award Number S10OD036363, the National Institute of General Medical Sciences of the National Institutes of Health [R35GM150765 (CPS), R35GM151249 (JMB), R35GM159898 (EH)]. The content is solely the responsibility of the authors and does not necessarily represent the official views of the National Institutes of Health. CPS also acknowledges the Wertheim UF-Scripps for start-up funds. The authors thank George Tsaprailis and Catherina Scharager Tapia at the UF-Scripps Proteomics Facility, and Robert M. Witwicki, Li Pan, and Marlene L. Biller at the UF-Scripps Genomics Facility.

## Author Contributions

CPS and EH conceived the work. CPS, EH, WZ, AL, CJD, JMB designed and executed the experiments. DWCM, EH, CPS directed research. CPS, WZ, AL, and EH prepared the manuscript.

## Author Information

The authors declare no competing financial interests. Readers are welcome to comment on the online version of the paper.

## Notes

### Competing Interest Statement

The authors have declared no competing interest.

