## Supporting information for "Mutant p53 binds RNA to drive mitochondrial dysfunction"

**Wuyue Zhou,[†,1,2] Alice Long,[†,3] Cameron J. Douglas,[1,2] Dalila Gallegos,[1] James M. Burke,[2,4] David W. C. MacMillan,[*3] Ezgi Hacisuleyman,[*2,4] Ciaran P. Seath[*1,2]**

[1]*Department of Chemistry, Wertheim UF Scripps, Jupiter, Florida 33458, USA.*

[2]*Skaggs Graduate School, The Scripps Research Institute, Jupiter, Florida, 33458, USA.*

[3]*Department of Chemistry, Princeton University, Princeton, NJ, USA.*

[4]*Department of Molecular Medicine, Wertheim UF Scripps, Jupiter, Florida 33458, USA.*

[†] Authors contributed equally.

### Supplementary Figures

#### Supporting Figure 1

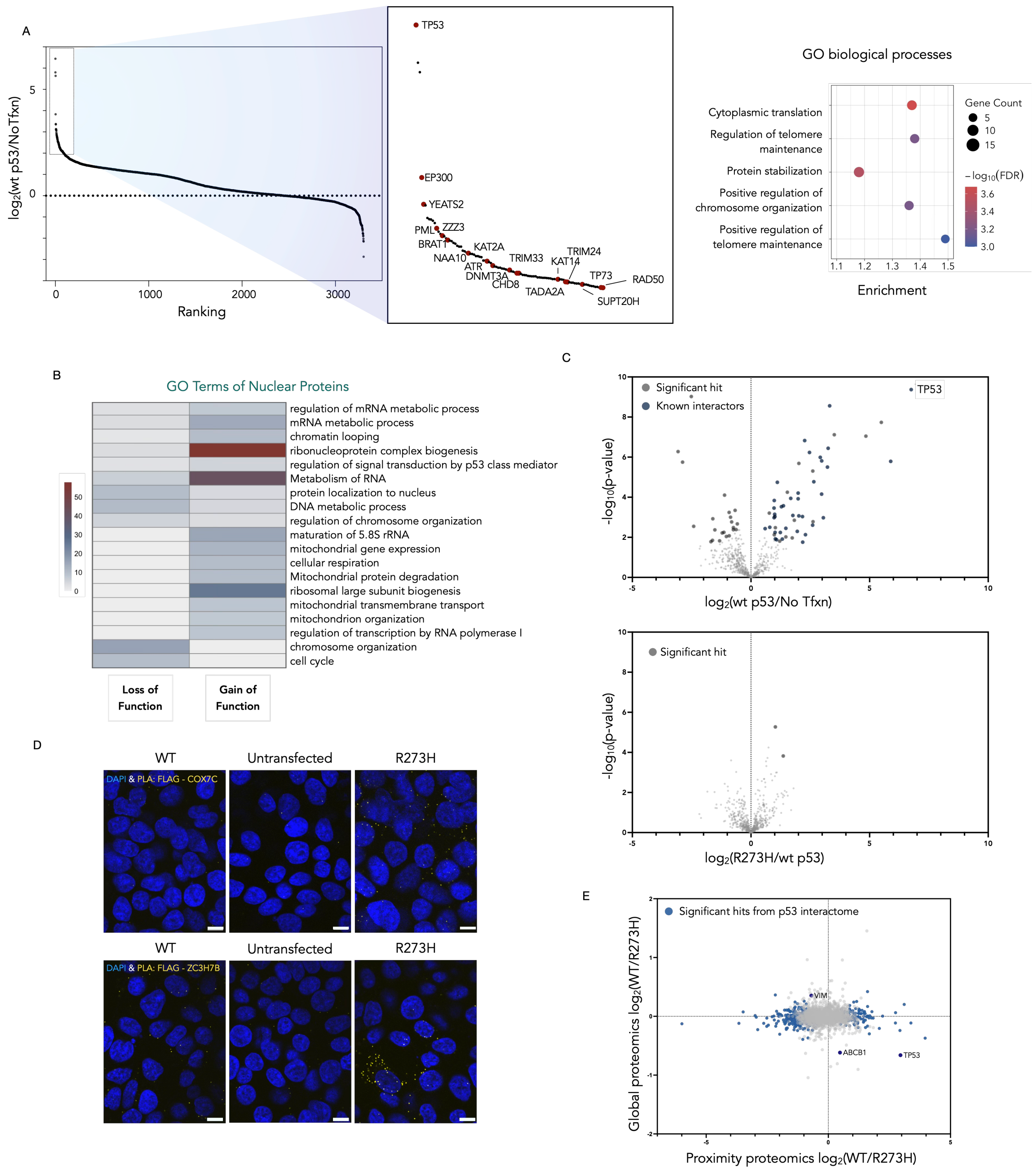

**Supplementary Fig.1 (A)** Intein-based μ-Map successfully revealed wt p53 interactome. Left: Waterfall plot with known p53 interactor annotated. Wild-type p53 was among the top hits. Right: GO biological processes. Terms are ranked by a weighted harmonic mean between the enrichment score and -$\log_{10}$FDR. **(B)** GO analysis of protein recruited to mutant R273H p53 (gain of function) and protein repulsed from mutant (loss of function). **(C)** IP-MS of p53. Top: A volcano plot derived from a two-sided t-test showing wt p53 interactors from IP-MS. Wild-type p53 was among the top hits, and known interactors were labeled. FDR<0.05. Bottom: A volcano plot derived from a two-sided t-test showing GoF and LoF of R273H p53 comparing to wt p53 from IP-MS. Few differential interactions were observed. **(D)** Representative images from PLA assay. Scale bar = 10 μm. **(E)** Scatter plots of μ-Map obtained p53 interactome shift (x-axis) and whole cell proteome (y-axis) changes in response to p53 mutation (R273H). Genes of interest are labeled and highlighted in blue.

### Supporting Figure 2

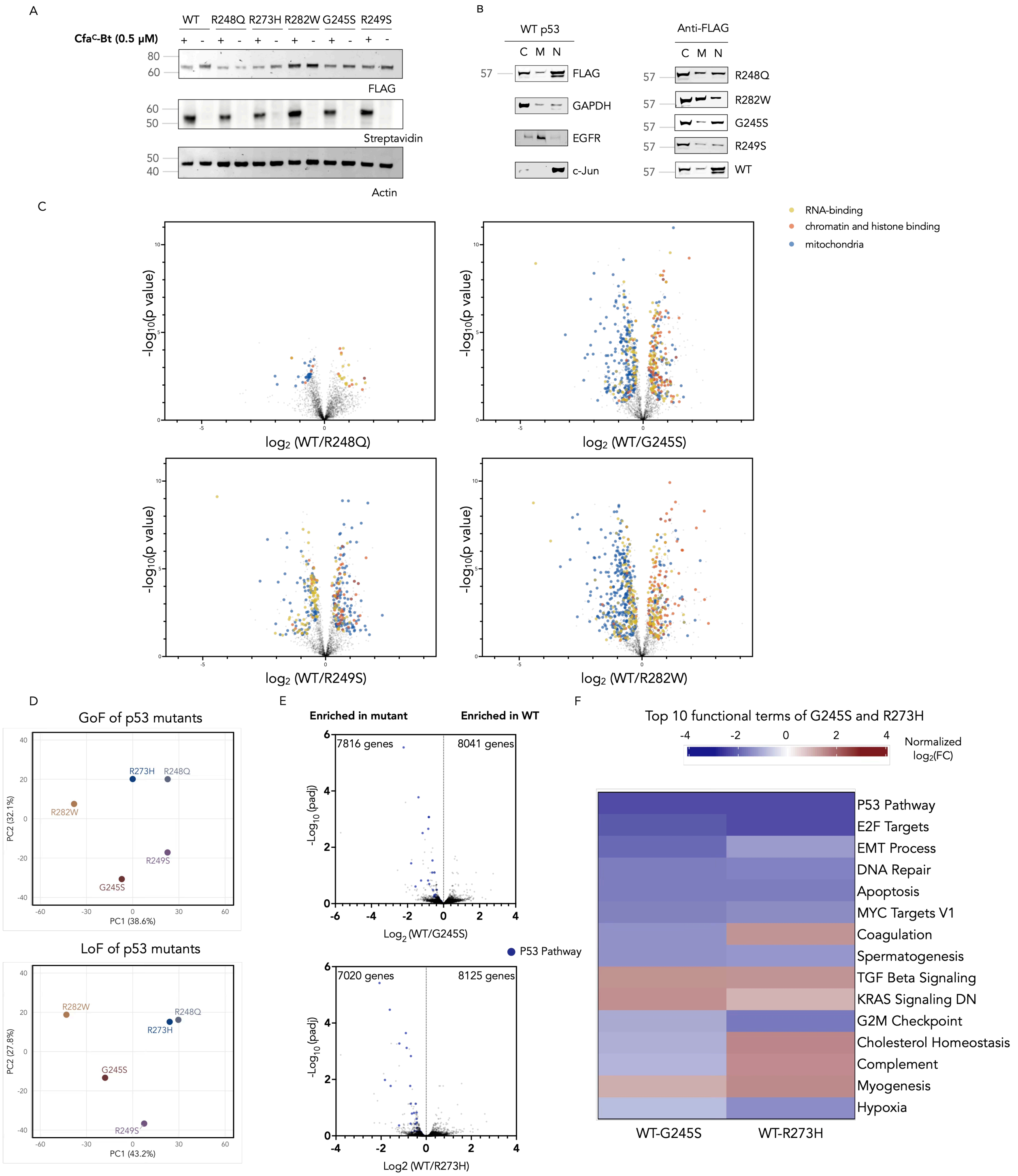

**Supplementary Fig.2 (A)** Spliced product is only observed while treated with Cfa$^{C}$-Bt for all p53 mutants used in this study. **(B)** Mutant and wt p53 distribution revealed by cellular fractionation. C= cytosol, M= cell membrane, and N= nucleus. Corresponding markers were also stained. **(C)** Interactome shift caused by p53 mutation. All volcano plots were derived from a two-sided t-test comparing wt p53 interactors to mutant p53 interactors. FDR<0.05. RNA-binding, chromatin and histone binding, and mitochondrial proteins are annotated. Right panel: enrichment of selected proteins in the dataset. **(D)** PCA analysis of gene annotations of p53 mutants. **(E)**

Transcriptome change caused by mutant p53 expression. Volcano plots were derived from a two-sided t-test comparing transcripts in wt p53 expressing cells to transcripts in mutant p53 expressing cells. Number of enriched genes are annotated. Genes relevant to p53 pathway are upregulated in both mutant forms of p53, and are annotated on the graph. **(F)** Combined top functional terms of differentially regulated genes in both mutants. Entries were ranked by adjusted p-values in ascending order.

### Supporting Figure 3

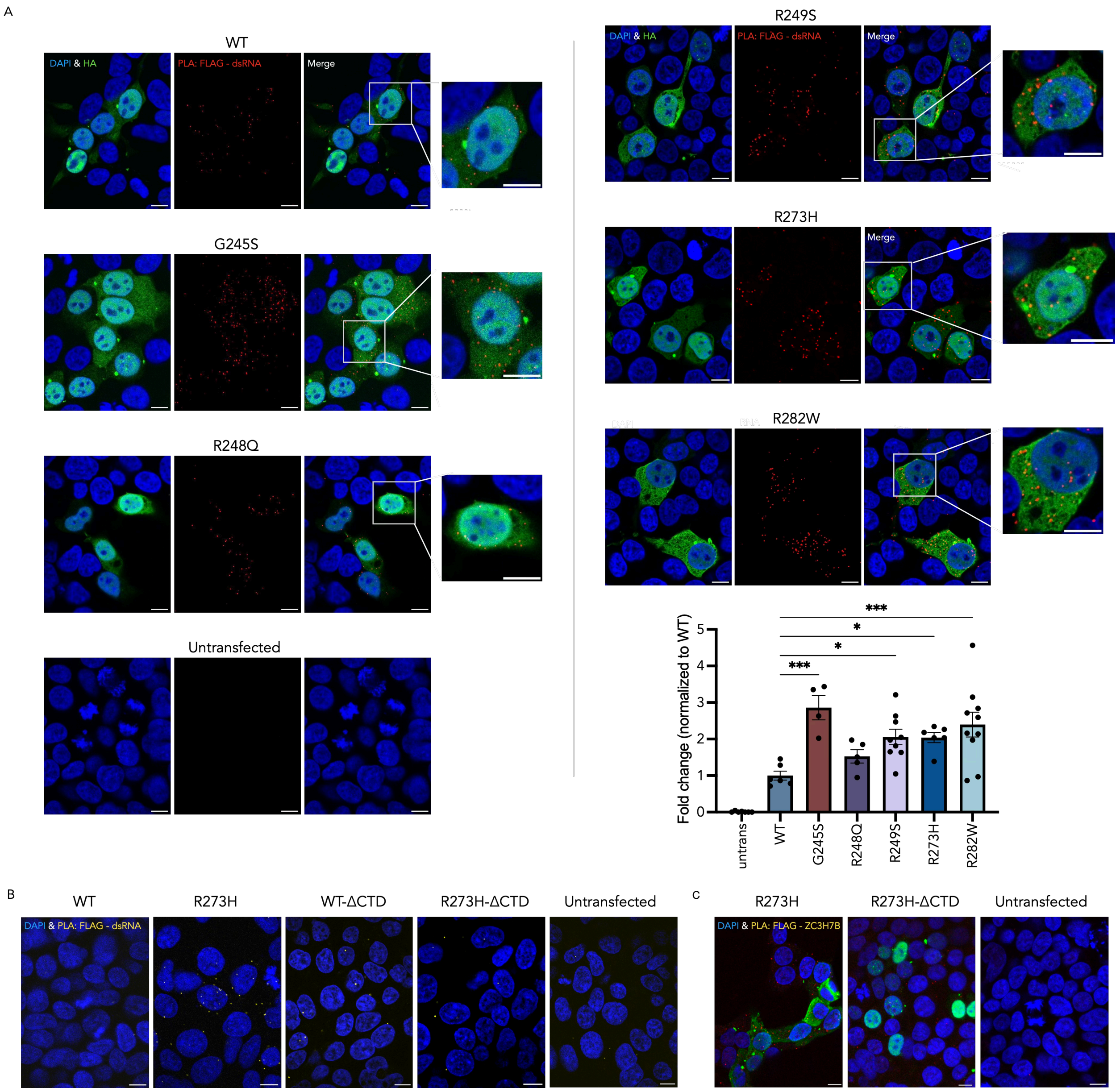

**Supplementary Fig.3 (A)** Representative images from PLA assay testing expressed p53 construct (FLAG) interaction with dsRNA (n ≥ 4, one-way ANOVA test followed by post hoc Dunnett's multiple comparisons comparing to untransfected control. *P<0.05, ***P<0.001). HA signal was stained to better localize transfected cells, scale bar = 10 μm. **(B)** Representative images from PLA assay testing expressed p53 construct (FLAG) interaction with dsRNA after CTD truncation, scale bar = 10 μm. **(C)** Representative images from PLA assay testing expressed p53 construct (FLAG) interaction with ZC3H7B, scale bar = 10 μm.

### Supporting Figure 4

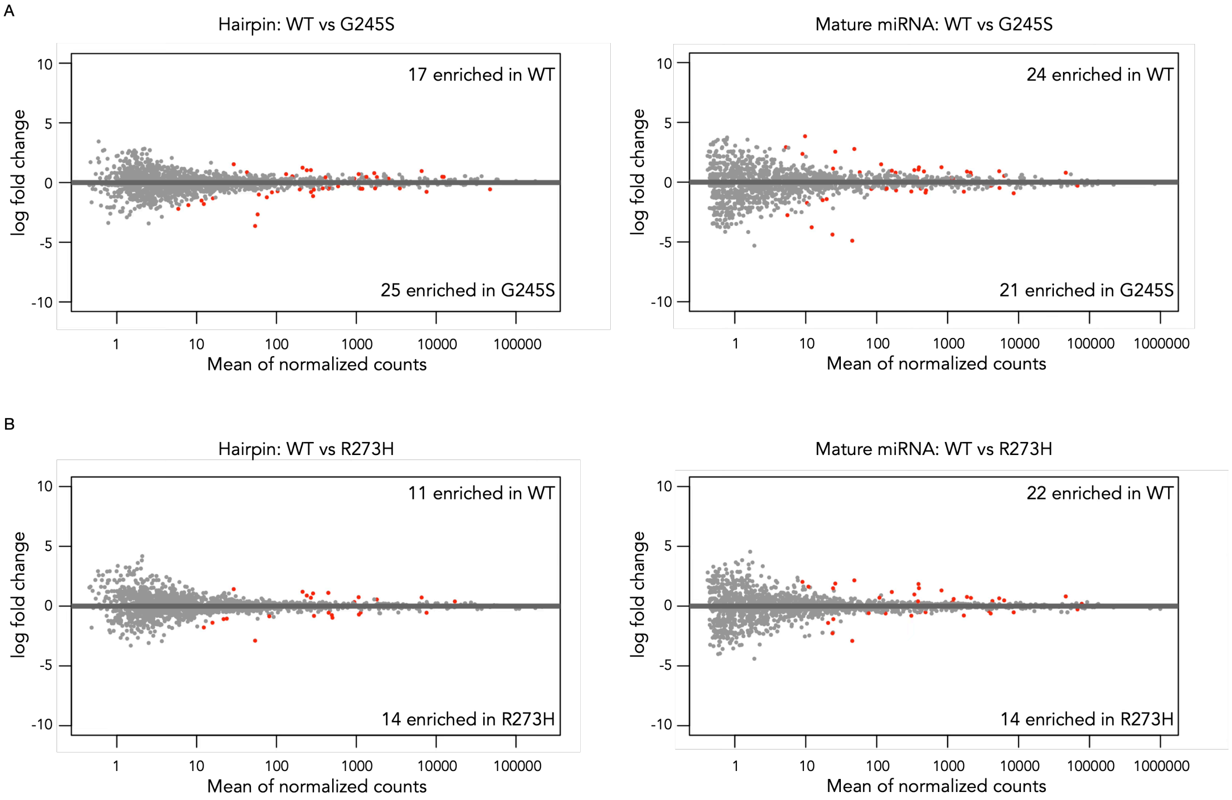

**Supplementary Fig.4 (A)** MA plot derived from DepSeq. Differentially regulated miRNA or hairpins in G245S mutant were highlighted. FDR<0.1. **(B)** MA plot derived from DepSeq. Differentially regulated miRNA or hairpins in R273H mutant were highlighted. Adjusted p-value<0.1.

### Supporting Figure 5

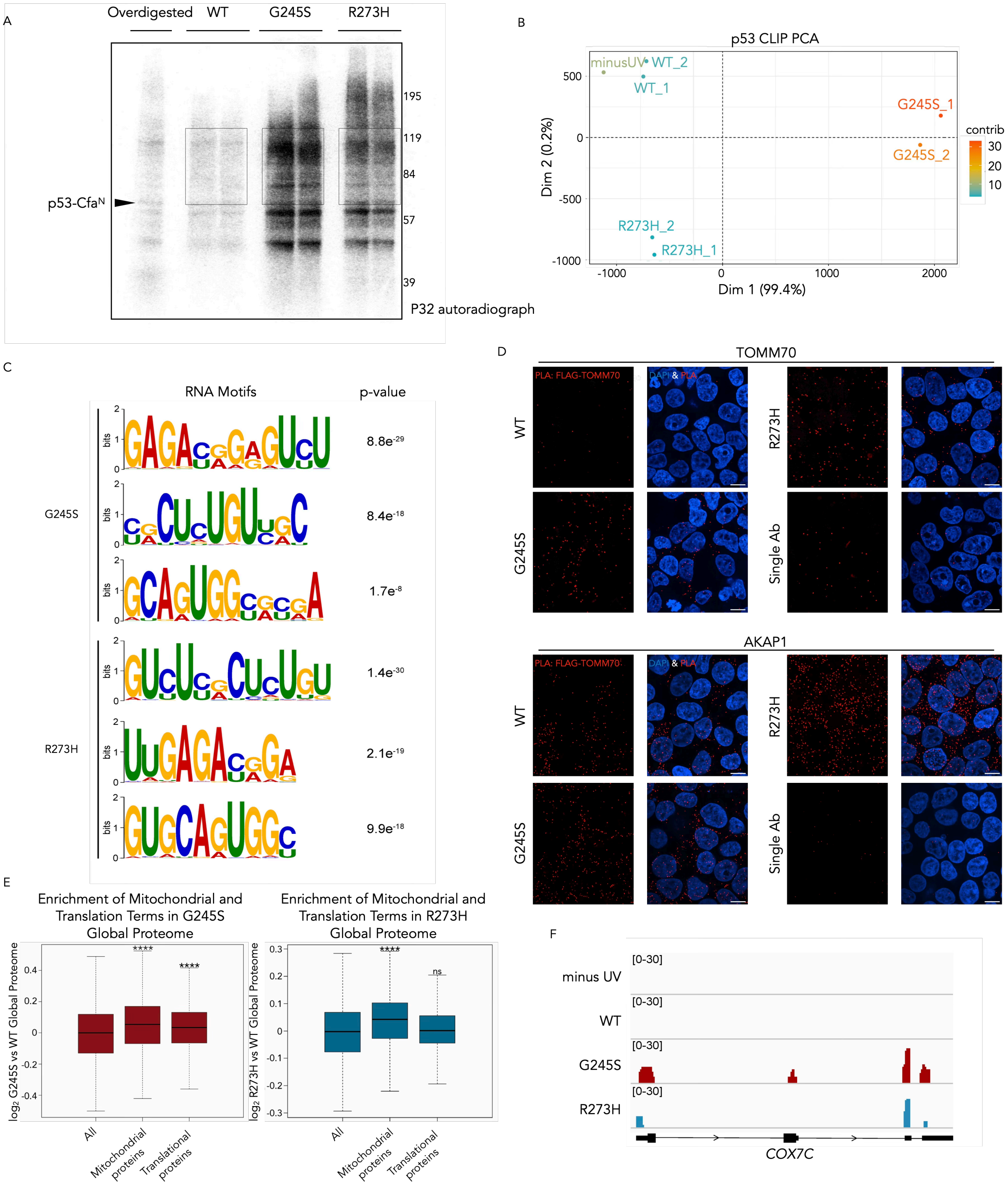

**Supplementary Fig.5 (A)** Representative autoradiograph image of CLIP-seq samples showing RNAs bound by wild type and mutant p53 after pull-down. Regions corresponding to p53 binding (labeled by dashed lines) were cut and subjected to digestion and purification to acquire RNA. **(B)** PCA plot showing CLIP-seq sample distribution. **(C)** Other consensus RNA-binding motifs of mutant p53 identified by STREME analysis. **(D)** Representative images from PLA assay testing expressed p53 construct (FLAG) interaction with TOMM70 or with AKAP1. **(E)** Among mutant p53 proteome, proteins associated with mitochondria (G245S and R273H) and translation (G245S) were highly enriched. Mitochondrial proteins were obtained from MitoCarta 3.0, and translation-related proteins were retrieved from MSigDB (GOBP: Translation; GO:0006412). Statistical significance was calculated using the Wilcoxon signed-rank test. **(F)** Visualization of CLIP-seq read coverage on transcript *COX7C*.

### Supporting Figure 6

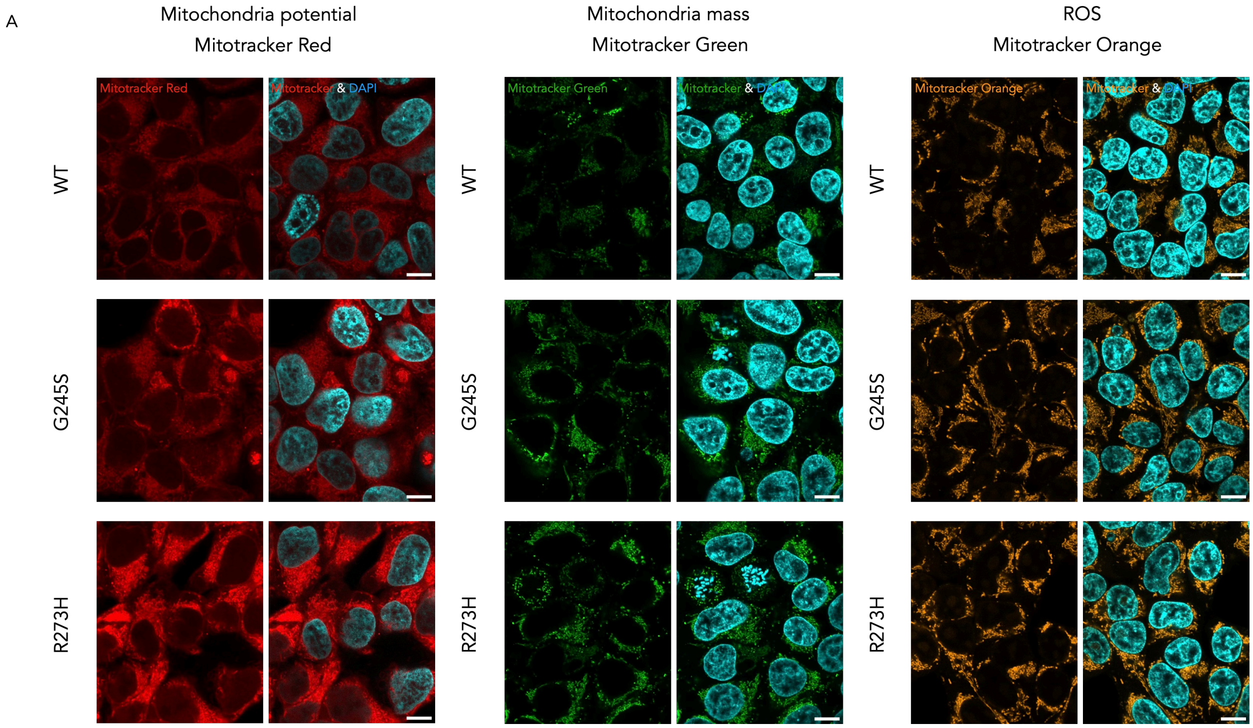

**Supplementary Fig.6 (A)** Representative images of Mitotracker assay. Scale bar = 10 μm.

### General Considerations

Water was purified using a Millipore Milli-Q Integral Water Purification System. All buffers and synthetic starting materials were used as received from commercial sources. Eppendorf Protein LoBind tubes (Z666505) were purchased from Millipore Sigma. RIPA Lysis buffer (10X, 20-188) was purchased from Millipore Sigma. Halt™ Protease Inhibitor Cocktail (100X, 78438), Pierce BCA Protein Assay Kit (23227), Bolt Bis-Tris Plus Gel (NW04120BOX) and iBright Prestained Protein ladder (LC5615) were purchased from Thermo Fisher Scientific. TBST (10X, TBST01-03) was purchased from Bioland Scientific LLC. DMEM (Gibco, 10566016), DPBS (Gibco, 14190250), Fetal Bovine Serum (Gibco, 10437-028), 100 U $mL^{-1}$ penicillin 100 μg $mL^{-1}$ streptomycin (Gibco 15140-122), and Trypsin-EDTA (Gibco, 25300054) were obtained from Thermo Fisher Scientific. Trypsin Protease (MS grade, 90057) was purchased from Thermo Fisher Scientific. Streptavidin Mag Sepharose magnetic beads (28985799) were purchased from Cytiva. Duolink® Proximity Ligation Assay kits (Duolink® In Situ PLA® Probe Anti-Rabbit PLUS, DUO92002; Duolink® In Situ PLA® Probe Anti-Mouse MINUS, DUO92004; Duolink® In Situ Detection Reagents Red, DUO92007) and Acetonitrile (271004) were purchased from Millipore Sigma.

#### Antibodies used in this study

Primary antibodies

| | | |
|---|---|---|
| Anti-FLAG (ms) | Cell Signaling (8146S) | 1:1,000 in TBST/ in antibody diluent |
| Anti-FLAG (rb) | Cell Signaling (14793S) | 1:1,000 in TBST/ in antibody diluent |
| Anti-FLAG (For IP) | Sigma Aldrich (F1804) | 8 µL per 100 µL Protein A Dynabeads |
| Anti-HA | Abcam (ab9110) | 1:1,000 in TBST |
| EGF Receptor | Cell Signaling (4267S) | 1:1,000 in TBST |
| c-JUN | Cell Signaling (9165S) | 1:1,000 in TBST |
| GAPDH | Cell Signaling (97166S) | 1:1,000 in TBST |
| p53(7F5) Rabbit | Cell Signaling (2527S) | 1:1,000 in TBST |
| ZC3H7B Poly Ab | Proteintech (25624-1-AP) | 1:1,000 in TBST/ in antibody diluent |
| dsRNA(K1) Mouse mAb | Cell signaling (28764S) | 1:1,000 in antibody diluent |

| | | |
|---|---|---|
| dsRNA(J2) Mouse Ab | Cell signaling (76651L) | 1:1,000 in antibody diluent |
| COX7C | Invitrogen (PA5-68018) | 1:1,000 in TBST/1:200 in antibody diluent |
| TOMM70 | Proteintech (14528-1-AP) | 1:500 in antibody diluent |
| AKAP1 | Proteintech (15618-1-AP) | 1:40 in antibody diluent |

Secondary antibodies

| | | |
|---|---|---|
| Goat Anti-Rabbit IgG H&L (Alexa Fluor® 488) | Abcam (ab150077) | 1:1000 in 2% BSA in PBST (IF)<br>1:5000 in 2% BSA in TBST (WB) |
| Goat Anti-Rabbit IgG H&L (Alexa Fluor® 555) | Abcam (ab150078) | 1:1000 in 2% BSA in PBST (IF)<br>1:5000 in 2% BSA in TBST (WB) |
| Goat Anti-Rabbit IgG H&L (Alexa Fluor® 647) | Abcam (ab150079) | 1:1000 in 2% BSA in PBST (IF)<br>1:5000 in 2% BSA in TBST (WB) |
| Goat Anti-Mouse IgG H&L (Alexa Fluor® 488) | Abcam (ab150113) | 1:1000 in 2% BSA in PBST (IF)<br>1:5000 in 2% BSA in TBST (WB) |
| Goat Anti-Mouse IgG H&L (Alexa Fluor® 555) | Abcam (ab150114) | 1:1000 in 2% BSA in PBST (IF)<br>1:5000 in 2% BSA in TBST (WB) |
| Goat Anti-Mouse IgG H&L (Alexa Fluor® 647) | Abcam (ab150115) | 1:1000 in 2% BSA in PBST (IF)<br>1:5000 in 2% BSA in TBST (WB) |
| Goat Anti Rabbit 680 (Li-Cor) | LICORbio (926-68071) | 1:5000 in 2% BSA in TBST |
| Goat Anti Rabbit 800 (Li-Cor) | LICORbio (926-32211) | 1:5000 in 2% BSA in TBST |
| Goat Anti Mouse 680 (Li-Cor) | LICORbio (926-68070) | 1:5000 in 2% BSA in TBST |
| Streptavidin IRDye 800CW (Li-Cor) | LICORbio (925-32230) | 1:5000 in 2% BSA in TBST |
| Goat Anti Rabbit IgG (H+L) Alexa 488 | Thermo Fisher (A32731) | 1:1000 in 2% BSA in PBST (IF)<br>1:5000 in 2% BSA in TBST (WB) |
| Goat Anti Mouse IgG (H+L) Alexa 488 | Thermo Fisher (A32723) | 1:1000 in 2% BSA in PBST (IF)<br>1:5000 in 2% BSA in TBST (WB) |

#### Cloning

To construct p53 plasmid conjugated to intein tags, pcDNA3.1(+)-H3.1-$Cfa^N$ plasmid (generous gift from Dr. D.W.C. MacMillan) were PCR cloned for backbone and intein expressing CDS using Phusion polymerase (New England Biolab (NEB), M0536S). The backbones were then fused with PCR amplified CDS region from commercial plasmid expressing wt p53 and ligated using In-Fusion snap assembly kit (Takara Bio, 638947). The encoded constructs were thus under the control of a CMV promoter. Ligated vectors were introduced to NEB 5-alpha competent E. coli cells (NEB, C2987H) by heat shock transformation. For mutant p53 expressing plasmids, mutagenesis was performed individually for every mutant using Q5® Site-Directed Mutagenesis Kit (NEB, E0554). For p53 proteins with CTD region deletion, deletion was performed with In-Fusion snap assembly kit. All primers were designed using SnapGene (version 7.2) and purchased from Integrated DNA Technologies (IDT). Ligated pcDNA3.1(+) vectors were introduced to NEB 5-alpha competent E. coli cells (NEB, C2987H) by heat shock transformation. The whole plasmid sequencing was performed by Plasmidsaurus using Oxford Nanopore Technology with custom analysis and annotation. For a detailed list of plasmids used in this study, refer to supplementary Table 1.

#### Cell culture

HEK 293T cells and stable HEK293T cells expressing mutant and wt p53 were cultured as a monolayer in DMEM, supplemented with 10% v/v FBS, 100 U $mL^{-1}$ penicillin, and 100 µg $mL^{-1}$ streptomycin. Cells were maintained in an incubator at 37 °C with 5% $CO_2$. For stable cell lines, media was supplied with 1.5 µg/mL Puromycin (InvivoGen, ant-pr-1).

#### Transient transfection

Each 10 cm plate of HEK293T cells at 50% confluency were transfected with a plasmid encoding mutant or wt p53 (5 µg per plate) with 15 µL of Lipofectamine 2000 (Thermo, 11668019) following the manufacturer's instructions. After 6 h, the media was aspirated and replaced with fresh media. Transfection was performed for 24 h in an incubator at 37 °C with 5% $CO_2$.

#### Generation of stable cell line

Concentrated lentivirus (>$10^8$ TU/mL) that expresses wt and mutant p53 were prepared by VectorBuilder. Briefly, to construct stable cell line, low passage HEK293T cells (24-well plates, 50% confluency) were incubated with optimal amount of lentivirus particles to achieve a MOI=4 for 24 h. Incubating media were supplied with 5 µg/mL of Polybrene (Millipore Sigma (Sigma),

TR-1003-G). Virus-containing medium was replaced with complete DMEM medium after 24 h and cells were incubated for another 24 h. Selective medium containing 1.5 μg/mL Puromycin (InvivoGen, ant-pr-1) were then replaced, and cells were maintained in the selective medium for 4 days to allow for complete selection.

#### RNA-seq analysis

Stable cells that express wt and mutant p53 from three different biological replicates were harvested, pelleted, and lysed with DNA/RNA Shield (Zymo Research, 1100-50) ($5 \times 10^5$ cells/ 50 μL). Lysed cells were submitted to Plasmidsaurus for RNA-seq analysis. In brief, FastQ was generated and demultiplexed with BCL Convert v4.3.6 and fqtk v0.3.1. Read-filtering was performed using FastP v0.24.0 with the following criteria: poly-X tail trimming, 3' quality-based tail trimming, a minimum Phred quality score of 15, and a minimum length requirement of 50 bp. Alignment to the appropriate reference genome was done using STAR aligner v2.7.11 with non-canonical splice junction removal and output of unmapped reads. Coordinate sorting of BAM files was performed using samtools v1.22.1. Removal of PCR and optical duplicates were done using UMICollapse v1.1.0. Alignment quality metrics, strand specificity, and read distribution across genomic features were performed using RSeQC v5.0.4 and Qualimap v2.3. Comprehensive QC report was generated using MultiQC v1.32. Gene-expression quantification was performed using featureCounts (subread package v2.1.1) with strand-specific counting, multi-mapping read fractional assignment, exons and three prime UTR as the feature identifiers, and grouped by gene_id. Final gene counts were annotated with gene biotype and other metadata extracted from the reference GTF file. Sample-sample correlations for sample-sample heatmap and PCA were calculated on normalized counts (TMM, trimmed mean of M-values) using Pearson correlation. Differential expression using edgeR v4.0.16 with filtering for low-expressed genes with edgeR::filterByExpr with default values. Functional enrichment performed for human and mouse samples using gene set enrichment analysis with GSEApy v0.12 using the MSigDB Hallmark gene set. Results were visualized with GraphPad Prism, and cross-analyzed with other omics datasets with R studio.

#### Label Free Global Proteomics

Stable cells expressing mutant or wt p53 from three independent biological replicates were lysed with SDS lysis buffer, and proteins were cleaned up, and digested using Sera-Mag Carboxylate SpeedBeads (Cytiva Life Sciences: E7 Cat. 45152105050250 and E3 Cat. 65152105050250) following manufacturer's protocol. Digested peptides were split into two technical replicates per sample.

Peptide digests were acidified with TFA to 0.1% (v:v) and desalted using 2 µg capacity ZipTips (Millipore, Billerica, MA) according to manufacturer instructions. Following drying under vacuum, peptides were re-solubilized in 0.1% formic acid (FA) to a final concentration of 100 ng/µL. Samples were analyzed on a nanoElute 2 (plug-in V2.1.79.0 (1); Bruker, Bremen, Germany) coupled to a Bruker TimsTOF Pro 2 mass spectrometer (Breman, Germany), equipped with a CaptiveSpray source and a 20µm zero dead volume (ZDV) Sprayer. Peptides (corresponding to 100 ng) were loaded onto a Thermo Fisher (Waltham, MA) PepMap Neo C18 trap column (300µm X 5mm, 5µm particle size) and then separated on an Aurora Series Elite reverse-phase C18 column (15cm X 150µm, 1.7µm particle size) from IonOpticks (Fitzroy, Australia). The column temperature was maintained at 50 °C using an integrated Bruker Column Toaster (Bremen, Germany). The column was equilibrated using 4 column volumes at 800 bar before loading samples in 100% buffer A (99.9% Fisher Optima® LC/MS water, 0.1% FA) at 201.6 bar. The trap column was equilibrated was equilibrated at 201.6 bar. Samples were separated at 500nL/min using a linear gradient from 2% to 35% buffer B (99.9% Fisher Optima® LC/MS acetonitrile, 0.1% FA) over 20.0 minutes before ramping to 95% buffer B (in 0.5 minutes) and sustained at 95% buffer B for 4.5 minutes (total separation method time 25.0 min). The Bruker TimsTOF Pro 2 was operated in DIA-PASEF mode using Tims Control v. 5.0.2. Settings for the MS method were as follows: Mass Range 100 to 1700m/z, 1/K0 Start 0.6 V.·/cm2 End 1.4 V·s/cm2, TIMS Ramp and accumulation time 75ms, Capillary Voltage 1700V, Dry Gas 3 l/min, Dry Temp 200°C, DIA-PASEF settings: 18 MS/MS scans (50m/z windows, 0.21 1/K0 windows, total cycle time 0.74), mass range 300 to 1200, and CID collision energy 20eV (at 0.60, 1/K0) to 65eV (at 1.60, 1/K0). The analysis was performed at The Herbert Wertheim UF Scripps Institute for Biomedical Innovation & Technology, Mass Spectrometry and Proteomics Core Facility (RRID:SCR_023576).

Data was processed via DIANN 1.8.1. Parameters set as follows: trypsin/P digestion, 3 missed cleavages, 3 max. variable modifications, N-term M excision, Ox(M), Ac(N-term) and C carbamidomethylation. Peptide length range was 7-30, precursor charge range 1-4, m/z range 300-1800, and fragment ion range 200-1800. Mass accuracy and MS accuracy were both set to 10. The following settings on the algorithm were checked: "Use isotopologues", "MBR", "No shared spectra", "Heuristic protein inference". Precursor FDR was set to 1%. A spectral library was used generated via DIANN from all known human proteins (In-Silico specral library). Resulting matrix.pg file was opened in Perseus (v2.0.7.0). Intensities were inputted as "main", the rest of the descriptors are categorical. Data was then transformed ($Log_2$). Data is annotated by treatment. Missing values were imputed with Perseus default settings. Normalization was performed via median subtraction. Following this process, a volcano plot was generated utilizing a t-test for statistical significance. The resulting volcano plots were plotted in GraphPad Prism 10 for final figures. Metascape and STRING was used for Gene Ontology.

#### General procedure for photoproximity labeling in cells

Peptides were prepared and purified as previously described previously.[1] $3\times10^7$ HEK 293T cells transfected with p53-HA-$Cfa^N$-FLAG were permeabilized by hypotonic pressure with 3 ml RSB buffer (10 mM tris, 15 mM NaCl, 1.5 mM $MgCl_2$, 1 × Halt protease cocktail, pH 7.6) for 10 min on ice. The cells were pelleted by centrifugation at 400 g for 5 min at 4 °C. The cells were resuspended in 3 ml RSB buffer, and homogenized with ten strokes of a loose pestle Dounce homogenizer, and pelleted at 400 g for 5 min at 4 °C. The cells were resuspended in crosslinking buffer (20 mM HEPES, 1.5 mM $MgCl_2$, 150 mM KCl, 1 × Halt protease cocktail, pH 7.6) and centrifuged at 400 g for 5 min at 4 °C. Finally, the cells were resuspended in 300 μL of crosslinking buffer per 1 $x10^7$ cells. To the suspended pellet was added $Cfa^C$-Ir in crosslinking buffer (0.5 μM final concentration). For splicing efficiency confirmation, $Cfa^C$-Bt was added in crosslinking buffer (0.25 μM final concentration). The cell suspension were incubated at 37 °C for 1 h.

The cells were isolated by centrifugation at 400 g for 5 min at 4 °C and washed twice with crosslinking buffer (500 μL) to remove excess peptide. The pellets were then resuspended in 3 mL crosslinking buffer containing diazirine-biotin conjugate (200 μM) and irradiated with blue light for between 60s in the Penn PhD Photoreactor M2 at 100% light intensity at 4 °C. The cells were re-isolated by centrifugation at 400 g for 5 min at 4 °C and washed once with crosslinking buffer to remove excess biotin-diazirine. The washed pellets were then resuspended in 2 mL LB3 buffer (10 mM tris, 100 mM NaCl, 1 mM EDTA, 0.5 mM EGTA, 0.1% sodium deoxycholate, 0.5% sodium lauroyl sarcosinate, pH 7.5) and sonicated using a Diagenode Bioruptor Plus for 12 cycles of 30 sec on and 30 sec off at high power at 4 °C. For splicing confirmation, pellets were collected and lysed as described immediately after the intein washing step and then subjected to western blot analysis.

The lysed cells were then clarified through centrifugation at 15,000 g for 20 mins at 4 °C and the protein concentration of the supernatant was determined by Pierce BCA assay. Protein concentration was normalized across all experimental replicates and diluted to 1 mg/mL with binding buffer (25 mM tris, 150 mM NaCl, 0.25% v/v NP-40, pH 7.5). 1 mL of each sample was then incubated with 100 μL of pre-washed magnetic Sepharose streptavidin beads for 2 h at RT with end-over-end rotation. The beads were subsequently washed twice with 1% w/v SDS in PBS, twice with 1 M NaCl in PBS, and 10% EtOH in PBS x 3. At this point proteins can be eluted from beads for western blotting by addition of 1 × Laemmli sample buffer (Biorad, 1610747) containing 20 mM DTT and 25 mM biotin and heated to 95 °C for 10 min.

#### Western blot analysis

Cell lysates were quantified with BCA and normalized to the same concentration and the cell lysate was boiled in 1X Laemmli buffer (diluted from 4x Laemmli Sample Buffer, supplied with 2.5 % v/v β-mercaptoethanol) at 95 °C for 10 min and loaded to a pre-cast Bolt™ 4-12% Bis-Tris Plus Mini Protein Gel (Thermo, NW04120BOX). Proteins were transferred to a nitrocellulose membrane (Thermo, 88018) and blocked with 2% Bovine Serum Albumin (BSA) (Fisher Scientific (Fisher), BP9704100) in TBST (diluted from 10x TBST (Bioland Scientific LLC, TBST01-03) with distilled water) for 1 h at RT. Membranes were then stained with primary antibodies diluted in 2% BSA in TBST at 4 °C overnight on a rocker. Membranes were then washed 3×10 min with TBST and stained with secondary fluorescent antibody diluted in 2% BSA in TBST for 1 h at RT. Membranes were then washed with TBST for 3x10 min and then imaged on Licor Odyssey CLx scanner. Quantification of the membrane was performed with ImageJ2 (2.14.0).

#### Label-free proteomics and data analysis of enriched sample

Following streptavidin-enrichment, the beads were resuspended in 0.5 mL PBS and transferred to a new tube. The beads were washed with 3 × 0.5 mL PBS and 3 × 0.5 mL 100 mM ammonium bicarbonate (Sigma, A6141). The beads were resuspended in 0.5 mL of 3 M urea (Sigma, U5378) in DPBS and 25 µL of 200 mM DTT in 25 mM ammonium bicarbonate was added. The beads were incubated at 55 °C for 30 min. Subsequently, 30 µL of 500 mM iodoacetamide in 25 mM ammonium bicarbonate was added and incubated for 30 min at room temperature in the dark. The supernatant was removed and the beads washed with 3x with 0.5 mL of DPBS and 6x with 0.5 mL of 50 mM ammonium bicarbonate. The beads were resuspended in 0.5 mL of 50 mM ammonium bicarbonate and transferred to a new tube. The beads were resuspended in 40 µL of 50 mM ammonium bicarbonate added with 1.2 µL of trypsin (1 mg/mL in 50 mM acetic acid (Fisher, A11350)), and incubated overnight with end-over-end rotation at 37 °C. After 16 hr, a further 0.8 µL of trypsin was added and the beads were incubated for an extra 1 h at 37 °C. Supernatant was then transferred to new tube and each biological replicate split into two technical replicates. Three biological replicates were used per condition. Peptides were treated and analyzed as described above.

#### Procedure for Immunoprecipitation MS

3×$10^7$ HEK 293T cells transfected with wt/mutant p53-HA-$Cfa^N$-FLAG grown in 15cm plates were washed 2-3x with ice-cold PBS, then lysed directly in the plate with EBC buffer (50mM Tris-HCl, 120mM NaCl, 0.1mM EDTA, 0.5% NP-40, 10% glycerol, 1× Halt protease inhibitor cocktail) and harvested via scraping. After lysing for 30 min on ice, the lysates were sonicated on the Diagenode Bioruptor Sonicator (LOW, 5 cycles at 4 °C, 7 s ON, 5 s OFF). Samples were then centrifuged (12,000 x g, 15 min, 4 °C), the supernatant was harvested, and all lysates were normalized to 1 mg/mL. 5 µg of the anti-FLAG antibody and isotype control antibody were then conjugated to Pierce Protein A Magnetic Beads (Thermo Scientific, 88845) according to the manufacturer's instructions. Then lysates were then incubated with the prepared protein A beads overnight at 4 °C with rotation. Beads were washed 4x with Wash buffer (50mM Tris-HCl, 150mM NaCl, 5mM EDTA, 0.1% Tween-20) and transferred to new tubes. Proteins were eluted with 1%

SDS in RIPA buffer at 95 °C for 10 min. Eluted protein was then precipitated on to speedbeads, washed and digested as described in the Label-free proteomics protocol above.

#### Procedure for cell fractionation

One 10 cm plate of HEK 293T cells was transfected with the p53 plasmids and cells were collected with trypsin. Cells were lysed in hypotonic lysis buffer (10 mM Tris, 15 mM NaCl, 1.5 mM MgCl2, PI, pH 7.6) on ice for 10 min, followed by centrifugation at 400g for 5 min at 4 °C. The supernatant was removed (cytosolic fraction) and the pellet was resuspended in hypotonic lysis buffer supplied with 1% v/v Triton X-100. Nuclei were lysed with ten strokes of a tight pestle homogenizer followed by centrifugation at 10,000 g for 10 min at 4 °C. The supernatant was removed (nucleoplasmic fraction) and RIPA buffer was added to the pellet. The sample was sheared by probe sonication (2 × 10 s total, 25% amplitude, 1 s on, 1 s off) to yield the chromatin fraction. SDS loading buffer was added and the samples were analyzed by western blotting using the indicated antibodies.

#### Immunofluorescence assay of p53-expressing cells

Stable cells or transiently transfected HEK293T cells that express p53 protein were plated on pre-treated round coverslips with poly-L-lysine (Sigma, P4707) in 24-well plates. When the cells reached 80% confluency (24 h post-transfection) they were gently washed with DPBS, then immediately fixed with 4% paraformaldehyde in PBS (Thermo, AAJ61899AP) for 10 min at RT and blocked and permeabilized for 1 h at RT with 2% BSA in PBST solution (1X DPBS, added with 0.1% Triton X-100 (Fisher, 501781842)). Cells were then stained with primary antibody diluted in 2% BSA in PBST solution at 4 °C ON. Following primary antibody staining, cells were washed with DPBS three times and stained with secondary fluorescent antibody for 1 h at RT. Coverslips were mounted with mounting media supplied with DAPI (Fisher, NC9524612) and sealed with clear nail polish on glass slides. Slides were imaged on an Olympus FV3000 laser scanning microscope with Abbelight system with a 100X UPLAPOHR OIL NA 1.5 WD 0.12 mm lens, with 405, 488, 561, and 640 nm laser. Colocalization signal analysis and line analysis was done via ImageJ2 and CellProfiler (version 4.2.8).

#### PLA analysis

Transiently transfected HEK293T cells that express p53 protein (for ZC3H7B, dsRNA, and COX7C), or stable cells that expresses p53 constructs (for TOMM70 and AKAP1 interaction) were plated on pre-treated round coverslips with poly-L-lysine in 24-well plates and fixed with 4% PFA in PBS 24 h post-transfection. Cells were incubated with pre-chilled 100% MeOH at -20 °C to permeabilize. Once fixed and permeabilized proximity ligation was carried out as recommended by manufacturer using the Duolink in situ red starter kit mouse/rabbit (Sigma-Aldrich DUO92101). For transient transfected cells, slides were stained with primary HA-tag antibody ON at 4 °C to identify p53-expressing cells, followed by 1 h of secondary antibody staining at RT. Slides were then mounted with duolink in situ mounting media with DAPI and sealed with clear nail polish for imaging. Slides were imaged on a FV3000 laser microscope as being described previously. At least 3 images were taken of each slide in different position. Number of PLA puncta signal were quantified using CellProfiler nuclear speckle counting software. Statistical significance was determined by a one-way ANOVA test followed by post hoc Dunnett's multiple comparisons test of foci per cell using GraphPad Prism.

#### miRNA-Seq of p53-expressing cells

HEK cells were seeded and transfected in 10 cm plates as described above. The next day, cells were harvested with trypsin, washed with PBS, and RNAs were extracted from $5x10^6$ cells using the RNeasy plus mini kit (Qiagen, 74134) according to the manufacturer's instructions. Total RNA was submitted to The Herbert Wertheim UF Scripps Institute for Biomedical Innovation & Technology - Genomics Core (RRID:SCR_017827), where it was quantified using a Qubit 2.0 Fluorometer (Invitrogen, Carlsbad, CA) and evaluated on an Agilent 4200 TapeStation (Agilent Technologies, Santa Clara, S10 CA) for quality assessment. All RNA samples had RNA Integrity Number (RIN) ≥7.0 and were used for total RNA-seq library preparation. RNase-free working environment was maintained, and RNase-free tips, tubes, and plates were utilized. 0.4 µg of total RNA per sample was used. The library preparation from the input RNA was conducted according to the NEXTFLEX small RNA kit v4 (Revvity, NOVA-5132-31). Briefly, the input RNA has been ligated to 3' adenylated adapter and 5' adapter. The RNA was then reverse transcribed to generate the first strand of cDNA. The synthesized product was cleaned up with beads and amplified with PCR (using 15 cycles) to incorporate a unique barcode and to generate final libraries. The libraries were purified using SPRI beads to remove any remaining primers and adaptors. The final libraries were validated on an Agilent 4200 TapeStation (Agilent Technologies, Santa Clara, CA), normalized to 4 nM, pooled equally, and loaded onto an Illumina NextSeq 2000 P3 300-cycle flow cell (Cat. 20040561, Illumina, San Diego, CA) at 750pM final concentration and sequenced using 2 x 155bp paired-end chemistry. On average, a 20.5 million reads pass filter were generated per sample. Raw and processed data files were uploaded to the NCBI Gene Expression Omnibus (GSE307813).

miRNA sequencing data were processed using the nf-core/smrnaseq pipeline[2] (version 2.4.0, doi: 10.1038/s41587-020-0439-x.) on the HiPerGator high-performance computing cluster with Nextflow (version 25.04.4) and Singularity (version 3.10.4). Raw FASTQ files were quality-checked with FastQC (version 0.12.1) and trimmed using cutadapt[3] (version 5.2) following the instruction of NEXTFLEX small RNA kit v4. Contamination was filtered with Bowtie2 (version 2.5.4). Reads were aligned to the miRbase mature miRNA and miRbase hairpin using Bowtie1 (version 1.3.1). Post-alignment processing of miRbase hairpin was done with SAMtools (version 1.2.1), edgeR (version 4.8.0), and mirtop (version 0.4.28). Alignment against human reference genome was performed with Bowtie1. MiRDeep2 (version 0.1.3) was used to perform novel and known miRNA discovery. Mirtrace (version 1.0.1) was adapted for miRNA quality control, and MultiQC (version 1.27.1) was used for raw read, alignment, and expression results present. Downstream analysis was performed with DESeq2 (v1.34.0) to identify differentially expressed (DE) genes. Low count miRNA was filtered out (total sum count in all replicates less than 10). Statistical significances were determined with pAdjusted $< 0.1$ for both miRNA and hairpin. Data was visualized using R studio.

#### Crosslinking immunoprecipitation (CLIP) and CLIP-seq analysis

Cells were washed twice with 1× PBS and UV-crosslinked on ice with one pulse of 400 mJ/cm² followed by one pulse of 200 mJ/cm². Cells were immediately scraped in fresh ice-cold 1× PBS and centrifuged at 5300 × *g* for 5 min at 4 °C. The resulting pellets were flash-frozen and stored at −80 °C. Two biological replicates were prepared, each from a single 15-cm dish. Cell pellets were resuspended in 1 mL of lysis buffer (1× PBS, 0.1% SDS, 0.5% sodium deoxycholate, 0.5% NP-40, supplemented with freshly added protease inhibitors), and immunoprecipitations were performed 2 h at 4 °C using a mouse monoclonal anti-FLAG antibody (8 µg conjugated to 100 µL of Protein A Dynabeads). Subsequent radiolabeling, RNA extraction, reverse transcription, and qPCR steps were performed as described previously[4] with minor modifications. During the protein digestion step, a 7 M urea incubation was introduced prior to RNA precipitation, as described by Ule et al.[5]. Wild-type and mutant samples were pooled after reverse transcription barcoding to increase yield and ensure uniform amplification. Libraries were sequenced on an Illumina NextSeq platform to generate 75-nt single-end reads.

Sequencing reads were processed using the CLIP Tool Kit (CTK)[6] following published guidelines[7]. Briefly, reads were quality-filtered, demultiplexed, trimmed to remove 5′ and 3′ linker sequences, and collapsed to eliminate PCR duplicates. Filtered reads were then aligned to the human genome (hg38) using BWA[8]. Mapped reads were further collapsed using fastq2collapse.pl from the CTK toolkit with default parameters. The genomic distribution of uniquely mapped CLIP reads was determined using bed2annotation.pl. Uniquely mapped CLIP tags from all samples were pooled, and peaks were identified using tag2peak.pl (CTK) with a significance cutoff of $p < 0.05$. For each sample, tag counts within significant peaks were quantified using the countOverlaps function in R. Differentially bound peaks between mutant and wild-type p53 were identified using DESeq, applying an adjusted $p$-value threshold of < 0.05.

#### RT-qPCR assay

For each independent biological replicate, $2x10^6$ cells were harvested with trypsin, washed with PBS, and RNAs were extracted using RNeasy plus mini. The RNA samples were treated with DNase (Invitrogen, AM1907) to remove genomic DNA. The RNA concentrations were measured using Qubit RNA BR Assay Kit (Thermo Scientific, Q10210) and normalized to the same concentration. cDNAs were synthesized with the SuperScript III First-Strand Synthesis System (Invitrogen, 18080051) according to the manufacturer’s protocol with random hexamers. The cDNA concentration was measured with Qubit ssDNA Assay Kit (Thermo Scientific, Q10212) and normalized to 5 ng/µL. 10 ng of cDNA was loaded in real-time qPCR reactions, which were performed in technical duplicates using the Power SYBR™ Green PCR Master Mix (Thermo Scientific, 4367659) and specified primer (COX7C: forward- CTTCCAGCAGCGGTATGTT reverse-AGCTAGTAACGACCACTTGTTT; GAPDH: forward-AATCCCATCACCATCTTCCAG reverse- CCTTCTCCATGGTGGTGAAGAC).

Quantification of gene expression was measured using the 5 Real-Time PCR System. Statistical analysis and data visualization was performed with GraphPad Prism.

#### Seahorse Assay

Mitochondrial respiration analysis (Seahorse assay) was performed at the Metabolic core at The Herbert Wertheim UF Scripps Institute for Biomedical Innovation. Per individual well in XF Pro Cell Culture Microplates (Agilent Technologies, 103792-100) pre-treated with poly-l-lysine, 2.5 $*10^4$ stable cells expressing p53 constructs were plated 16 h prior to assay performing. To assess mitochondrial respiratory function, in whole cells, the growth media was replaced with XF media (unbuffered DMEM, pH 7.4, Agilent Technologies, 103680-100) with 10 mM pyruvate and 2.5 mM glucose supplied. The cell plate was equilibrated for 30 min at 37 °C in a $CO_2$-free incubator before being loaded in the XF-96 analyzer. Inside the analyzer, the plate was combined with the pre-calibrated XF cartridge with the individual well ports A, B and C loaded with 10× concentrated stocks of the following compounds, respectively: A. Oligomycin (20 µg/mL); B. FCCP (20 µM); C. Rotenone and Antimycin A (20 µM each). Oxygen consumption rate (OCR) was monitored in real-time at 37 °C under basal conditions and following the sequential delivery of the compounds in the cartridge ports. Non-mitochondrial rotenone/antimycin-insensitive OCR was subtracted from all other OCR measurements.

#### Mitotracker labeling

Per individual well in 12-well glass bottom plate (0.17 mm) (Cellvis, P12-1.5H-N), $3*10^5$ stable cells expressing p53 constructs were plated 16 h prior to assay performing. Cells were treated with warm media supplied with MitoTracker at 37 °C for 30 min. A final concentration of 250 nM MitoTracker Red (Thermo, M22426), 100 nM MitoTracker Green (Thermo, M7514), or 100 nM MitoTracker Orange (Thermo, M7511) were used. Hoechst dye (Thermo, H3570, 1 µg/mL) was added during the final 10 min of incubation time. Cells were immediately imaged on an Olympus FV3000 laser scanning microscope with Abbelight system with a 100X UPLAPOHR OIL NA 1.5 WD 0.12 mm lens, with 405, 488, 561, and 640 nm laser. At least 3 images were taken from two independent biological replicates. Signal analysis was done via ImageJ2. Images were transformed to 8-bit, thresholding was performed with Default ignore black white method, and intensity was measured for quantification. Statistical significance was determined by a one-way ANOVA test followed by post hoc Dunnett's multiple comparisons test of foci per cell using GraphPad Prism.
